# Cell-type specific subtyping of epigenomes improves prognostic stratification of cancer

**DOI:** 10.1101/2024.11.27.625781

**Authors:** Qi Luo, Andrew E. Teschendorff

## Abstract

**Background:** Most molecular classifications of cancer are based on bulk-tissue profiles that measure an average over many distinct cell-types. As such, cancer subtypes inferred from transcriptomic or epigenetic data are strongly influenced by cell-type composition and do not necessarily reflect subtypes defined by cell-type specific cancer-associated alterations, which could lead to suboptimal cancer classifications.

**Methods:** To address this problem, we here propose the novel concept of cell-type specific combinatorial clustering (CELTYC), which aims to group cancer samples by the molecular alterations they display in specific cell-types. We illustrate this concept in the context of DNA methylation data of liver and kidney cancer, deriving in each case novel cancer subtypes and assessing their prognostic relevance against current state-of-the-art prognostic models.

**Results:** In both liver and kidney cancer, we reveal improved cell-type specific prognostic models, not discoverable using standard methods. In the case of kidney cancer, we show how combinatorial indexing of epithelial and immune-cell clusters define improved prognostic models driven by synergy of high mitotic age and altered cytokine signaling. We validate the improved prognostic models in independent datasets, and identify underlying cytokine-immune-cell signatures driving poor outcome.

**Conclusions:** In summary, cell-type specific combinatorial clustering is a valuable strategy to help dissect and improve current prognostic classifications of cancer in terms of the underlying cell-type specific epigenetic and transcriptomic alterations.

## Introduction

The Cancer Genome Atlas (TCGA) ^1^ has transformed our molecular understanding of cancer and proposed many novel clinically relevant cancer classifications ^2–7^. These cancer taxonomies have, by and large, been derived from omic profiles generated in bulk-tissue, encompassing mixtures of many different cell-types. Whereas classifications based on somatic mutations and copy-number alterations reflect the underlying patterns of genomic alterations in tumor cells, classifications derived from bulk transcriptomic and epigenetic data are subject to potential confounding by cell-type heterogeneity (CTH) ^8–10^. Indeed, it is now well-known that inter-individual variation in tumor-tissue composition is substantial ^9–12^ and that this can strongly influence tumor classification ^9,13–15^. Although these classifications have often been shown to be of prognostic and clinical relevance (e.g. immune-reactive vs immune-cold tumors) ^2,16^, it is important to note that much of this inter-individual variation in cell-type composition (e.g. immune-cell infiltration) is also present within tissues of a healthy population ^11,12,17^. At the epigenetic level, obtained cancer classifications often reflect cell of origin ^9,18^, but are not necessarily informative of the epigenetic changes in the tumor cell-of-origin. In general, CTH implies that it is much harder to pinpoint whether specific cancer-associated transcriptomic or epigenetic changes are happening in the tumor cells and not in tumor stroma. It follows that most of the transcriptomic and epigenetic cancer taxonomies proposed to date do not necessarily reflect cancer subtypes defined by underlying cell-type specific molecular changes. The need to explore novel cancer classifications in terms of their cell-type specific transcriptomic and epigenetic changes is critical for an improved understanding of how distinct cancer subtypes emerge in relation to the functional changes that happen in tumor cells and in the various types of tumor-associated stromal cells.

Whilst single-cell technologies, notably scRNA-Seq ^19^, snRNA-Seq ^20^ and scATAC-Seq ^21^ are yielding novel insights into the tumor-stroma interface at cell-type resolution ^22–26^, as well as refining existing tumor classifications ^15^, single-cell studies on their own are limited to profiling relatively small numbers of tumor samples, which prevents us from capturing the extensive inter-subject clinical heterogeneity that we know exists. Indeed, at present, small-scale single-cell studies need to be combined with large-scale bulk-tissue datasets like the TCGA/ICGC in order to refine cancer taxonomies, as shown recently in the context of colorectal cancer ^14,15^ or breast cancer ^27–29^. Moreover, for certain data-types such as DNA methylation, profiling of single-cells, even in very modest numbers of clinical samples, is still not feasible ^30–32^. Thus, here we propose a novel strategy, based on the concept of “cell-type specific combinatorial clustering” (CELTYC), to refine the molecular classification of cancer-types. The key innovative idea behind this proposal is to perform clustering over the features (CpGs/genes) driving cancer-relevant cell-type specific variation, which results in cancer subtypes that reflect the changes in individual cell-types, and which are hence not merely driven by variations in cell-type composition. We evaluate the above CELTYC strategy in the context of DNA methylation (DNAm) data, which is less noisy than mRNA expression, allowing for more accurate estimation of cell-type fractions and cell-type deconvolution. The ability to estimate cell-type fractions with a reasonably high accuracy is indeed a critical step in our proposed strategy. By applying CELTYC to liver hepatocellular and kidney renal cell carcinoma, we reveal novel biologically and clinically relevant tumor classifications that significantly outperform existing ones in terms of associations with clinical outcome, highlighting the importance of cell-type specific subtyping of cancer.

## Results

### Cell-type specific clustering to refine molecular classifications of cancer

We reasoned that if molecular profiles representing clinical samples are clustered without adjustment for the underlying cell-type heterogeneity, that this could lead to suboptimal or skewed classifications of disease that are overly influenced by sample variations in cell-type composition. To demonstrate this, we first devised a simulation model where we mixed together sorted immune-cell DNAm profiles (**Methods, SI fig. S1a**). The mixtures were generated using Illumina 450k DNAm data from BLUEPRINT ^33^, representing sorted monocytes, neutrophils and CD4+ T-cells for each of 139 individuals. Cell-type proportions within mixtures were chosen from realistic estimates obtained in a tissue like blood, which we note results in significant variation in a given cell-type fraction across individuals (**Methods, SI fig. S1a**). The simulation model further assumes that samples consist of two disease subtypes distinguished by differential DNAm at a well-defined set of CpGs, but only in one immune-cell type (monocytes) (**Methods, SI fig. S1a**). We note that because in practice the sample subtype would be unknown, no supervised analysis is possible. With this simulation model, we thus asked if unsupervised clustering of the samples over the most variable CpGs would reveal the two molecular subtypes or not? SVD/PCA analysis clearly shows that the top PCs do not correlate with disease status, and a scatterplot of the top 2 PCs did not reveal segregation of samples by disease status (**SI fig. S1b-c**). Instead, mixtures segregate according to the neutrophil fraction, as evidenced by a near-perfect correlation of PC1 with this cellular fraction (**SI fig. S1c**). Clustering of the mixtures over the 1000 most variable CpGs also did not result in clusters associated with disease status (**SI fig. S1c**). Thus, this highlights how CTH can mask putative disease-relevant subtypes. Although this analysis only considered sorted immune cell-types, similar considerations would apply to solid tissues, where variations in underlying cell-type fractions are also substantial ^12^. To see this, we applied our EpiSCORE DNAm-atlas ^34^ to 15 TCGA cancer-types to estimate cell-type fractions in all tumor samples, revealing, in each cancer-type, that top PCs correlate most strongly with variations in underlying cell-type fractions (**SI fig. S2**). This suggests that current TCGA transcriptomic and epigenetic classifications are strongly influenced by variations in cell-type composition, potentially masking other, clinically more relevant, cancer subtypes.

Thus, to address this CTH challenge, we devised a computational strategy called CELTYC (Cell-Type Specific Combinatorial Clustering) (**Methods**, **Fig. 1**). Given a data matrix defined over CpGs and samples, CELTYC begins by estimating cell-type fractions in each sample using an algorithm such as EpiDISH (if the tissue is blood) ^35,36^, or the relevant tissue-specific DNAm reference matrix from the EpiSCORE DNAm-atlas ^37,38^ (for solid tissue-types) (**Fig. 1a**). These estimated fractions are then regressed out of the data matrix to construct the standardized residual variation matrix, reflecting data variation that is not caused by variations in cell-type composition (**Methods, Fig. 1b**). CELTYC then proceeds by running the CellDMC algorithm ^39^ to identify cancer-associated differentially methylated cytosines in each cell-type (DMCTs) (**Fig. 1c**). CellDMC uses statistical interaction terms between the phenotype (i.e. normal/cancer status) and cell-type fractions to infer cancer-associated DNAm changes in each cell-type ^39^. Typically, this results in a partitioning of DMCTs into a group of DMCTs that is common to all cell-types, other groups where DMCTs are only shared between specific cell-types, and finally groups of DMCTs that only appear in one specific cell-type (**Fig. 1c**). These partitions define separate data matrices, which can then be analyzed together or in combination using a statistical procedure called JIVE (Joint and Individual Variation Explained) ^40,41^, that extracts out components of joint variation across all DMCT submatrices as well as components of individual variation that are unique to each cell-type or unique to specific combinations of cell-types (**Fig. 1d**). Individual variation matrices represent unique variation only present in one cell-type which can be further analyzed with unsupervised clustering for novel class discovery (**Fig. 1e**). Alternatively, having inferred DMCTs in each cell-type, one can perform an unsupervised clustering of the standardized residual matrix as defined over the DMCTs of the given cell-type (**Fig. 1f**). The unsupervised clustering can thus reveal molecular subtypes defined by the DNAm changes in one particular cell-type, which could be very distinct from the subtypes identified by unsupervised clustering over bulk-tissue. Potentially, this could lead to improved prognostic models (**Fig. 1g**).

**Figure 1:**
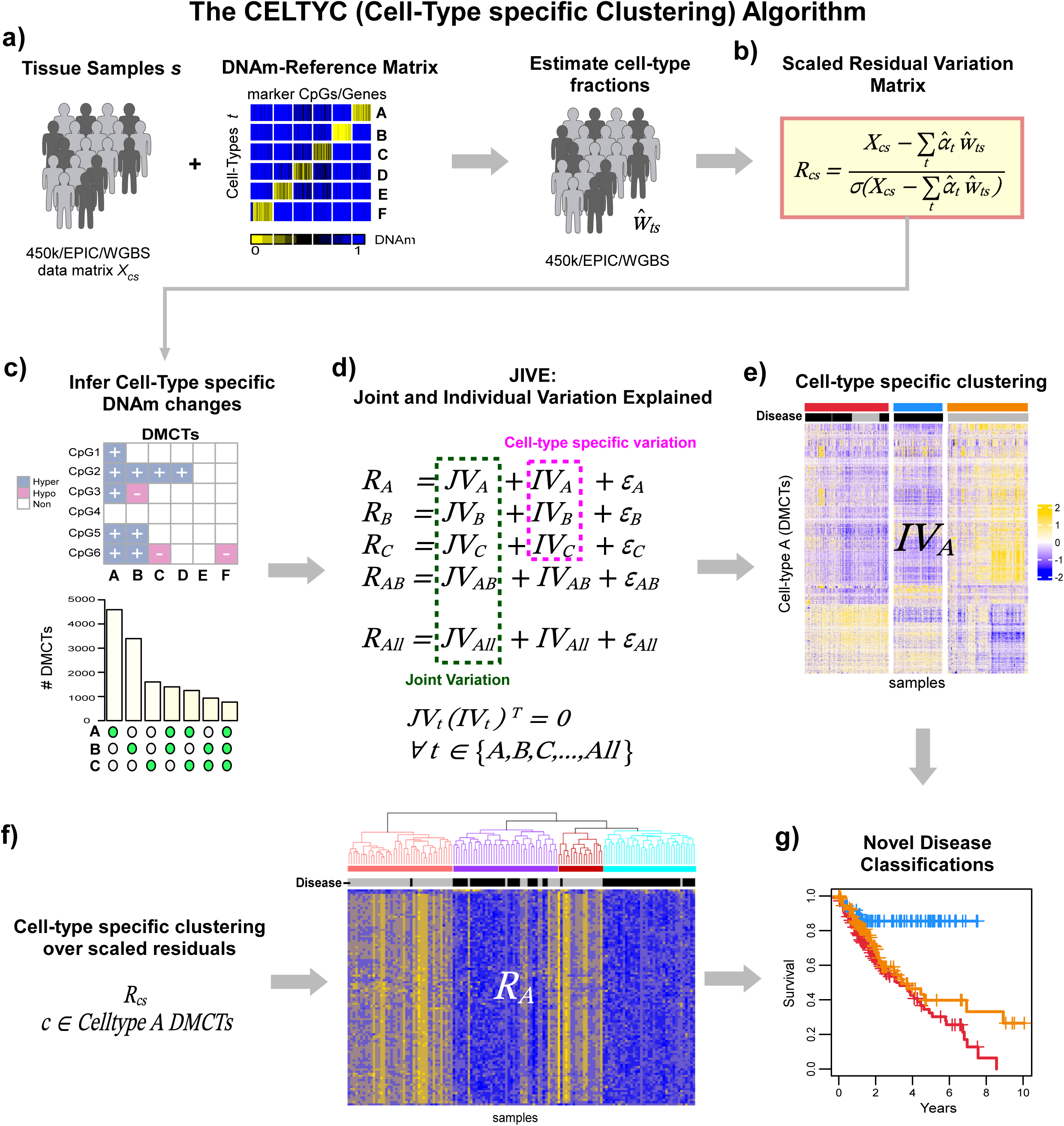
The CELTYC algorithm. **a)** Given a DNAm data matrix *X_cs_* for a collection of tissue samples *s* from a given tissue-type, we take a corresponding tissue-specific DNAm reference matrix defined over the main cell-types in the tissue to estimate cell-type fractions w…_ts_ for each cell-type *t* in sample *s.* **b)** From these cell-type fractions, we can generate the scaled residual variation matrix *R_cs_* over features (genes/CpGs) *c* and samples *s.* **c)** From this residual variation matrix we can then infer cell-type specific DNAm changes (DMCTs) in relation to some phenotype of interest, for instance using the CellDMC algorithm. **d)** We then perform JIVE clustering over residual matrices defined over mutually exclusive sets of DMCTs that may include the cell-type specific DMCTs and those shared by specific cell-types. JIVE extracts components of variation common to all the sets of DMCTs and unique to each set of DMCT. **e)** One could then cluster over the individual variation matrix defined for one cell-type (say “A”) to infer novel subclasses in terms of the DNAm changes that occur in that cell-type. **f)** Alternatively, one may cluster directly over the residual variation matrix defined over a specific set of DMCTs. **g)** Both e) and f) can lead to identification of novel prognostically relevant disease subtypes.

### Validation of CELTYC and power calculation on simulated data

To validate CELTYC, we first considered a simulation model in lung tissue. We simulated 108 bulk lung tissue samples by mixing together 108 bronchial epithelial cell (BEC) Illumina 450k DNAm samples from Magnaye et al ^42^ with 139 sorted monocyte, 139 CD4+ T-cell and 139 neutrophil 450k DNAm samples from BLUEPRINT ^33^ (**Methods, Fig. 2a**). To simulate disease-associated DNAm changes, we used the top 1000 differentially methylated cytosines (FDR<0.05) derived by comparing the BECs of 71 asthma cases to 37 non-asthmatic controls ^42^. Although these DMCs are related to a specific disease (asthma), their effect sizes are small and representative of those found in many other diseases, including cancer. The weights of the mixtures representing cell-type fractions were derived from realistic estimates inferred by applying EpiSCORE ^34^ to the large eGTEX lung-tissue DNAm dataset ^43^ (**Methods, SI fig. S3a**). Principal Component Analysis over the artificial mixtures confirmed that the top-PCs only correlated with variations in cell-type fractions and not with case/control status (**Fig. 2b**). Clustering over the most variable CpGs also did not reveal any segregation of samples by disease status (**SI fig. S3b**). This, once again, highlights the need to adjust for CTH. Hence, we first estimated the cell-type fractions of the simulated mixtures using our lung DNAm reference matrix (**Methods**), which revealed excellent agreement with the true (known) fractions (**SI fig. S3c**). Because on this dataset, the full conditional CellDMC model lacks power, we applied the marginal unconditional variant of CellDMC to infer disease-associated DMCTs **(Methods**). This revealed that most of the DMCTs occurred in BECs, as required, and that we could recover nearly 50% of the previously declared 1000 ground-truth disease-DMCs (**Fig. 2c**). Consequently, clustering the scaled residual variation matrix over these disease BEC-DMCTs revealed clear segregation by disease-status (**SI fig. S3d**). Of note, ordinary DMC-analysis without adjustment for CTH would only have detected a very small fraction of disease-associated DMCs (**Fig. 2c**). Next, we extended the previous simulation model to perform a power calculation that would inform us on the required sample size for the CELTYC strategy to work. We devised a parametric resampling scheme to simulate larger datasets of cases and controls (**Methods**), revealing the need to consider datasets encompassing at least 400 samples, in order to achieve significant sensitivity under the full conditional CellDMC model (**Fig. 2d**).

**Figure 2:**
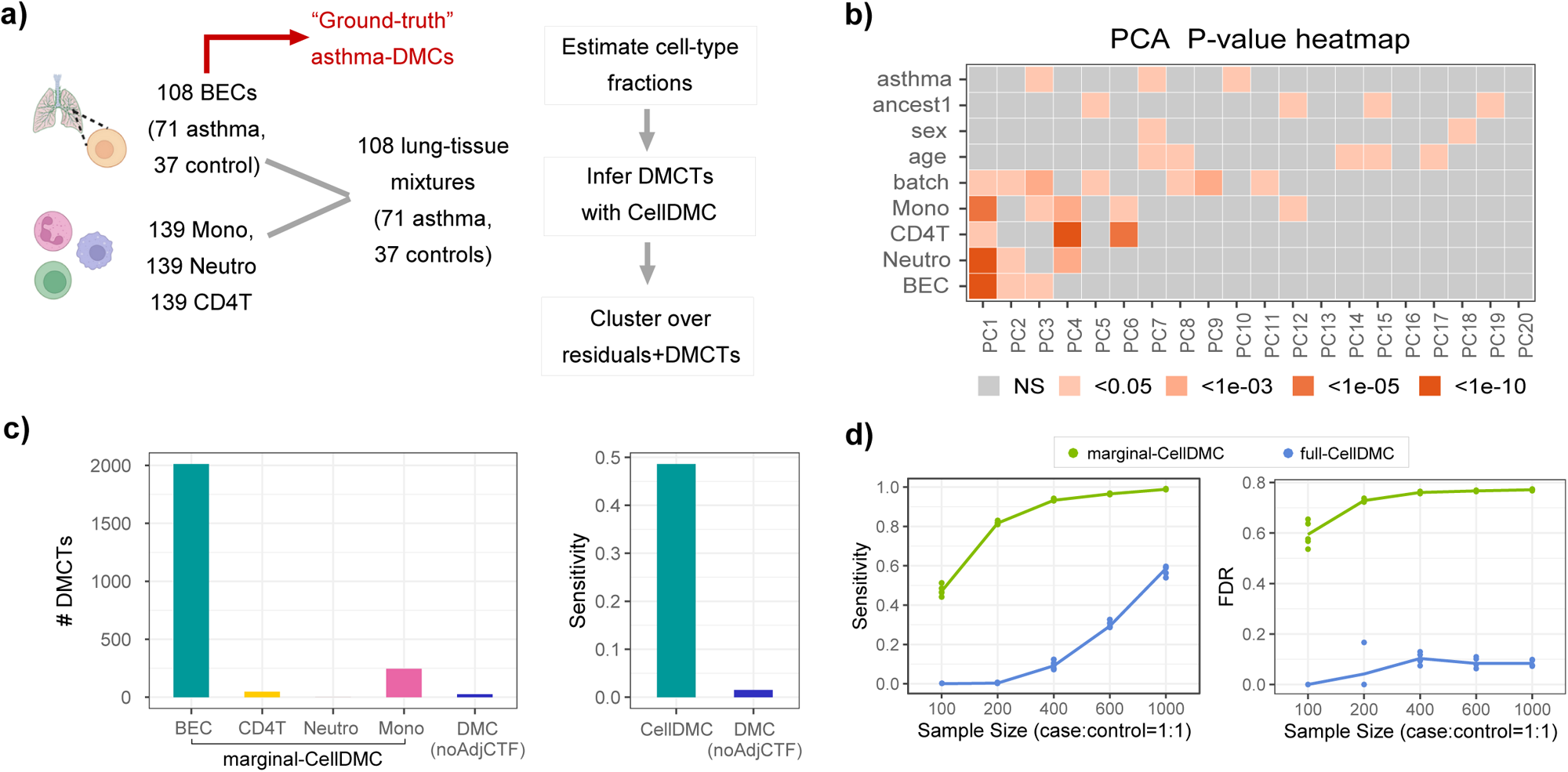
Validation of CELTYC on simulated data and power-analysis. **a)** Scheme of simulation model where Illumina 450k DNAm profiles of 108 sorted bronchial epithelial cells (BECs), 139 sorted neutrophils, monocytes and CD4+ T-cells from BLUEPRINT were mixed together using realistic cell-type fractions for lung-tissue as estimated from the eGTEX lung DNAm dataset consisting of over 200 lung tissue-samples. The BEC samples were from adult children with (cases) and without asthma (controls). Thus, the 108 lung mixtures are also from cases and controls. Cell-type fractions in the 108 mixtures were estimated using HEpiDISH, and subsequently CellDMC with these fractions was applied to infer DMCTs associated with case/control status. Sensitivity to capture the “ground-truth” disease-associated DMCs in BECs was estimated. Finally, clustering over the scaled residual variation matrix and BEC-DMCTs is performed. **b)** Heatmap of P-values of associations between principal components (PCs) and various factors, with the PCA performed on the 108 lung mixtures. **c)** Left: Number of disease-associated DMCTs (y-axis) in each cell-type as inferred by applying the marginal CellDMC model to the 108 lung mixtures. DMC labels the number of DMCs inferred using limma without adjustment for cell-type heterogeneity. Right: Sensitivity to capture the ground-truth disease-associated DMCs, defined as the top-ranked 1000 DMCs by comparing asthma BECs to non-asthma BECs. DMC labels the sensitivity using limma without adjustment for cell-type heterogeneity. **d)** Left: Sensitivity to detect the 1000 ground-truth disease-associated DMCs for simulated lung mixtures of different sizes (5 Monte-Carlo runs at each sample size), using the full conditional and marginal CellDMC models. Of note, in this simulation we assumed an equal ratio of cases and controls, so that a sample size of 200 means 100 cases and 100 controls. Right: as left, but for the false discovery rate (FDR).

### CELTYC identifies prognostic hepatocellular carcinoma subtypes

Based on the above power calculation, we thus applied CELTYC to cancer-types with sufficient numbers of cases and controls and to corresponding tissues for which reasonably accurate cell-type fractions can be estimated. We first considered the liver hepatocellular carcinoma Illumina 450k DNAm dataset (LIHC, 50 normal-adjacent samples, 379 cancers) from the TCGA ^44^, because for liver-tissue we had previously validated a DNAm reference matrix defined over 5 cell-types (hepatocytes, cholangiocytes, endothelial cells, Kupffer macrophages and lymphocytes) ^34^. Applying EpiSCORE ^37^ with this liver DNAm reference matrix, we estimated fractions for the 5 liver cell-types in all TCGA LIHC samples (**Fig. 3a**). We then applied CellDMC ^39^ to identify cancer-associated DMCTs (**Fig. 3a, SI table S1**). Most changes were observed in lymphocytes (n=4591), hepatocytes (n=3394) and endothelial cells (n=1602), with substantial overlaps between them (**Fig. 3b**). The number of cell-type specific DMCTs (i.e. those not shared with any other cell-type) was also largest for lymphocytes and hepatocytes, whilst the number of DMCTs unique to endothelial cells was much reduced (**SI fig. S4a**). Next, we applied consensus clustering ^45^ to the standardized residual variation matrix defined over lymphocyte-DMCTs, and separately also for hepatocyte and endothelial-DMCTs, revealing inferred clusters that were broadly consistent between cell-types (**SI fig. S4b**). Effect sizes between clusters, as defined in the original unscaled basis, were typically in the range of 1-30% DNAm change for the hepatocyte-DMCT defined clusters, but generally much smaller for lymphocyte and endothelial-DMCT ones (**SI fig.S5**). Importantly, the major inferred cell clusters for each cell-type displayed significantly different clinical outcome (**SI fig.S4c**). To see which cell-type may be driving the association with outcome, we repeated the clustering analysis but now restricting to cell-type specific DMCTs (i.e. upon removing the overlapping DMCTs) (**Fig.3c-d**), which revealed differences in clinical outcome only when clustering over lymphocyte-specific DMCTs (**Fig.3e**), suggesting that lymphocyte-DMCTs drive the classification patterns in relation to LIHC prognosis. Very similar results were obtained had we used JIVE to decompose the residual variation matrices into joint and cell-type specific components (**SI fig.S6**).

**Figure 3:**
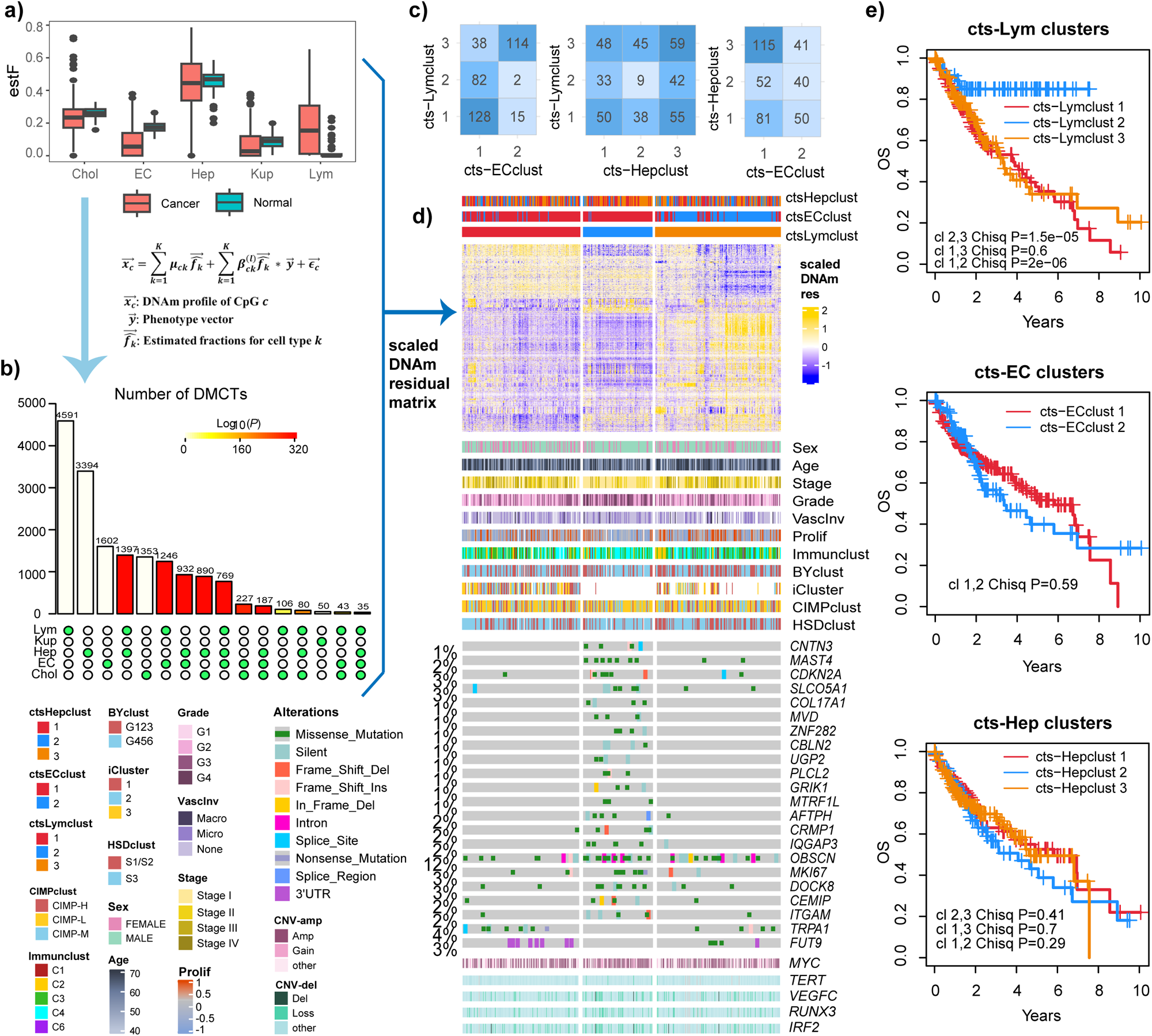
CELTYC identifies novel clinically relevant subtypes in liver hepatocellular carcinoma (LIHC). **a)** Boxplot of estimated fractions of cholangiocytes (Chol), endothelial cells (EC), hepatocytes (Hep), Kupffer cells (Kup) and Lymphocytes (Lym) in the normal and cancer LIHC TCGA DNAm dataset, as estimated using EpiSCORE. The estimated fractions are then used in linear interaction terms with normal-cancer status to infer cell-type specific differentially methylated cytosines (DMCTs) using the CellDMC algorithm. **b)** Landscape plot depicting the number of DMCTs in each cell-type as well as the numbers that overlap between cell-types. P-values of overlap were computed using the SuperExactTest. **c)** Confusion matrices for consensus clusters obtained with CELTYC using lymphocyte specific DMCTs, endothelial specific DMCTs and hepatocyte specific DMCTs. **d)** Clustering heatmap of the standardized/scaled residual DNAm variation matrix, with samples grouped by the 3 inferred consensus clusters obtained using lymphocyte specific DMCTs (ctsLym). On the top of the heatmap we also display sample labels representing the main clusters obtained using only endothelial specific DMCTs (ctsEC) or hepatocyte specific DMCTs (ctsHep). Below heatmap we display sample labels for sex, age, grade, stage, vascular invasion, proliferation index and other LIHC classifications (including Immune, Boyault (BY), iCluster, CIMP-based and Hoshida (HSD)), as well as somatic mutational, and copy-number profiles for key driver genes in LIHC. **e)** Kaplan Meier curves for the clusters inferred using CELTYC on lymphocyte specific DMCTs, hepatocyte specific DMCTs and endothelial cell (EC) specific DMCTs. P-values derive from a one-tailed Chi-Square test.

Reflecting differences in clinical outcome, the good prognosis LC2 cluster (i.e. cluster-2 of lymphocyte-DMCT specific clusters in **Fig. 3e**) displayed lower proliferation (P=0.018) and was predominantly low stage (stage 1+2) (P=1e-6) compared to the other two main clusters (LC1+LC3) (**Fig.3d**). On the other hand, LC2 was predominantly high grade (P=2e-7, **Fig.3d**). Whilst this may be surprising, it is worth noting that among all potential prognostic factors (i.e. age, sex, vascular invasion, tumor grade, stage, proliferation index and the novel proposed LC clusters), only stage (P=1e-6), proliferation index (P=4e-6) and LC clusters (P=5e-6) were significantly correlated with clinical outcome according to a univariate Cox proportional hazard regression, with all these associations remaining significant in a multivariate Cox model including all three variables (**SI table S2**). Thus, although the LC-classification from CELTYC correlates with proliferation and stage, it is also prognostically independent of these factors.

### CELTYC improves prognostic stratification of LIHC

Clustering over lymphocyte-specific DMCTs led to an excellent prognostic stratification with LC2 displaying an 80% overall survival rate 8 years after diagnosis, with LC1+3 displaying a corresponding survival rate of less than 20% (**Fig. 4a**). Thus, we next asked if this classification could have been obtained by more standard means. The importance of including lymphocyte-specific DMCTs was evident because using all other DMCTs resulted in poor discrimination accuracy (**Fig. 4b**). To assess the importance of CELTYC, we re-clustered LIHC samples over cancer differentially methylated positions (DMPs, FDR < 0.05) adjusted for cell-type fractions, which did not result in clusters displaying different clinical outcome (**Fig. 4c**). Clustering LIHC samples over the 1000 most variable CpGs also did not lead to good prognostic separability (**Fig. 4d**). Thus, these data indicate that CELTYC is a critical element in driving the strong prognostic separability. Next, we asked if CELTYC’s model improves upon state-of-the-art prognostic models for LIHC. We compared prognostic stratification of CELTYC to previous LIHC classifications, including the integrative iClusters from the TCGA ^44^, the immune-cluster subtypes from Thorsson et al ^16^, the CpG-island methylation phenotype subclasses ^46^, Hoshida’s mRNA-expression based classification ^47^ and Boyault’s subtypes ^48^ (**Methods**). Since the Hoshida, Boyault and CIMP classifications were not available, we computationally reproduced them (**Methods, SI fig.S7**). The prognostic separability of all these previous classifications was lower than the one obtained with CELTYC (**Fig. 4e-i**). Consistent with this, we note that in general there was no strong overlap between our CELTYC classification and these previous ones (**Fig.3d, SI table S3, SI fig.S8**). To formally prove that CELTYC’s prognostic model outperforms all other ones, we used a comparative likelihood-based strategy (**Methods**), which confirmed that CELTYC defines an improved prognostic model (**Fig. 4j**). Collectively, these results indicate that clustering over cell-type specific DMCTs, which avoids confounding by variations in cell-type composition, can reveal novel and improved prognostic subtypes, otherwise hidden by such variation. To facilitate future applications of CELTYC’s prognostic model, we used 10-fold cross-validation to derive a logistic Elastic Net DNAm-based predictor for the poor and good outcome CELTYC clusters (**SI table S4, Methods**), achieving a significant hazard ratio (HR=1.78, P=5e-06) in the cross-validation folds.

**Figure 4:**
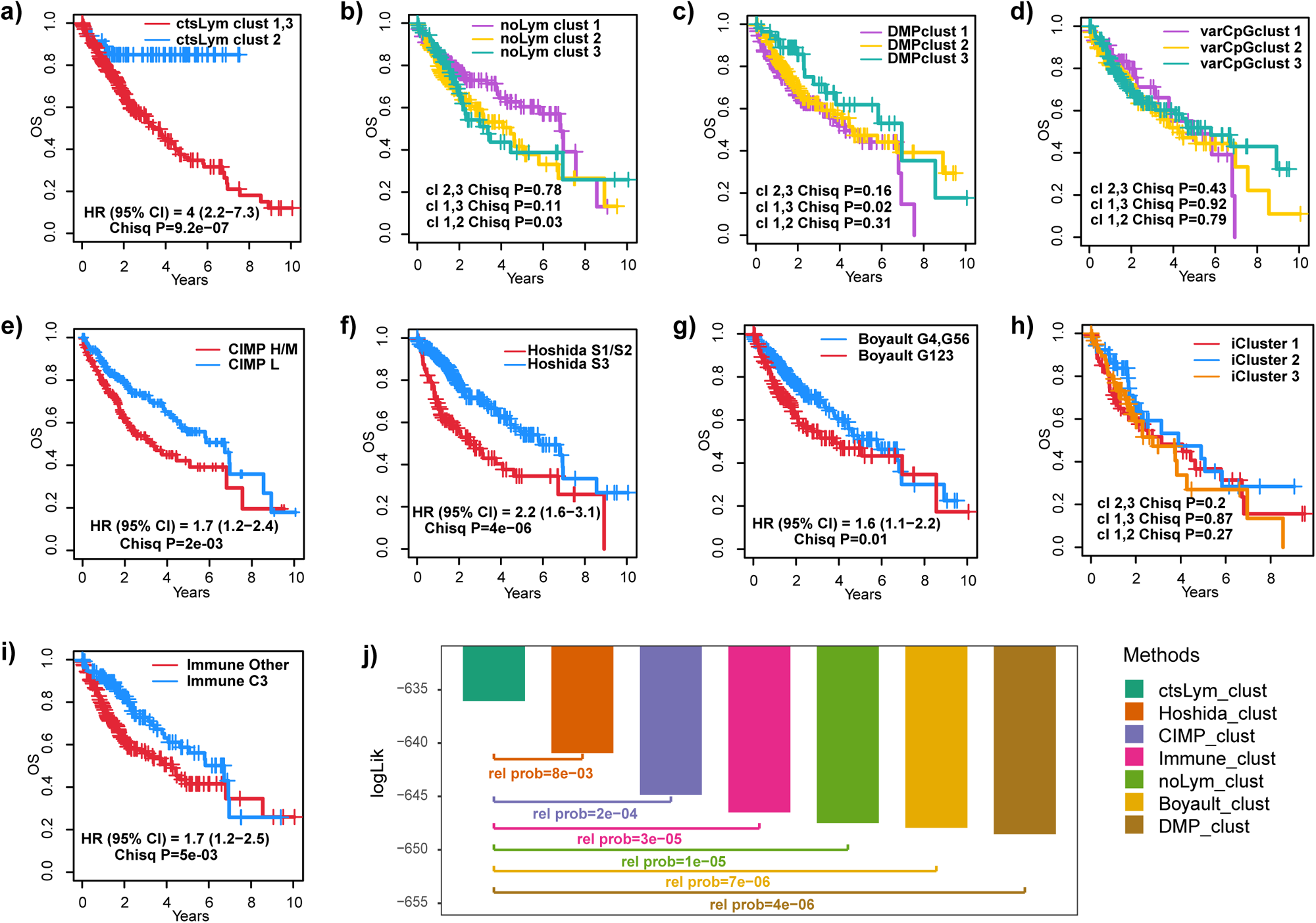
CELTYC improves prognostic separability of LIHC. **a)** Kaplan Meier curves for overall survival (OS) for the LC2 and LC1+3 clusters obtained by applying CELTYC to the lymphocyte specific DMCTs. Hazard Ratio (HR), 95% confidence interval and Chi-square test P-value is given. **b)** As **a)**, but for the three consensus clusters obtained if we use all DMCTs for all cell types except lymphocyte DMCTs. **c)** As **a)** but for the consensus clusters obtained by clustering over differentially methylated probes (DMPs, FDR<0.05 adjusted for cell-type fractions) between normal and cancer. **d)** As **a)**, but for the consensus clusters obtained by clustering over the 1000 most variable CpGs. **e)** Kaplan Meier curves for the CIMP-based clusters. **f)** Kaplan Meier curves for the Hoshida-clusters. **g)** Kaplan Meier curves for the Boyault clusters as shown. **h)** Kaplan Meier curves for the 3 TCGA iClusters. **i)** Kaplan Meier curves for the immune clusters. **j)** Barplot depicts the log-likelihoods of Cox-regressions of the different prognostic models with overall survival as endpoint, using the ordinal clusters of the models as predictors. In the barplot, we also display the relative probability of alternative prognostic models (i.e. CIMP, Hoshida etc) being better than the ctsLym-model. Red dashed line refers to a relative probability of 0.05, so any value less than this means that the ctsLym-model has a >95% chance of being a better prognostic model.

### CELTYC’s prognostic subtypes validate in independent LIHC datasets

Although CELTYC’s prognostic classification was derived by unsupervised clustering over cancer-DMCTs and thus unlikely to represent a false positive finding, we nevertheless aimed to validate this prognostic classification in independent LIHC datasets. Due to lack of availability of other large LIHC DNAm datasets, we decided to validate the prognostic subtypes in LIHC mRNA expression data. To this end, we first applied a logistic lasso classifier on the TCGA LIHC mRNA expression samples to predict their good and poor outcome CELTYC assignments, using a 10-fold CV-procedure to avoid overfitting (**Methods**, **SI fig.S9a**, **SI table S5**). We then applied this lasso-predictor to two independent LIHC mRNA expression datasets encompassing 100 and 247 hepatocellular carcinoma samples, respectively ^49,50^ (**Methods**). In both cohorts, the predicted poor and good outcome groups displayed significantly different overall survival (**SI fig.S9b, SI table S6**), with a prognostic separability similar to that observed in the TCGA cohort. Thus, this shows that the CELTYC prognostic model generalizes to independent cohorts and that it can be defined at the mRNA expression level. Interestingly, CELTYC’s prognostic model, which is inherently unsupervised, performed similarly to a supervised Elastic Net Cox-regression model trained against overall survival on the TCGA data (**SI table S7**).

### CELTYC’s classification is independent of somatic mutations and CNVs

Having validated CELTYC’s prognostic model, we next asked how CELTYC’s classification relates to somatic mutations and copy-number variations (CNVs) (**Methods**). We identified 178 genes whose mutation frequency differed between good and poor outcome groups (Fisher Exact Test P < 0.05), with the overwhelming majority (n=176) displaying a lower mutation frequency in the poor outcome class. Only 2 genes (*FUT9* and *TRPA1*) displayed more alterations in the poor outcome group (**Fig.3d**), and for only 4 genes (*FUT9, MUC6, MARVELD2, FER1L6*) did the somatic mutational profile directly correlate with clinical outcome (**SI table S8**). We verified that CELTYC’s prognostic model correlated with clinical outcome independently of these mutations, and that it also defined a stronger prognostic model than any mutational profile (**SI table S8**). A total of 2796 genes displayed a significant difference in gain or deletion/loss frequency between good (LC2) and poor outcome (LC1+3) subtypes (Fisher Exact Test P < 0.05), with the overwhelming majority displaying higher frequency of CNV-changes in the good outcome group. For instance, 1497 genes displayed a higher frequency of deletion or loss in the good outcome group, including LIHC tumor suppressors *RUNX3* ^51^ and *IRF2* ^52^ (**Fig.3d**). Among the genes with significantly different CNV frequencies between good and poor outcome clusters, a total of 618 genes were significantly associated with clinical outcome in univariate Cox-regression analysis, yet the CELTYC classification remained strongly prognostic when adjusting for any one of these (**SI fig.S10, SI table S9).** Overall, these results indicate that CELTYC’s classification is independent of somatic mutational or CNV-profiles.

### Epigenetic dysregulation of WNT signaling in poor outcome cluster

To shed light on the biological nature of CELTYC’s prognostic model, we first performed eFORGE analysis ^53,54^ of cell-type specific cancer-DMCTs stratified by hyper vs hypomethylation. Although this revealed some specificity, for instance, hypermethylated DMCTs in cancer hepatocytes and hypomethylated DMCTs in cancer endothelial cells were enriched for the active TSS chromatin state as defined in liver and endothelial cells, respectively (**Fig. 5a**), this enrichment analysis was insufficient to shed light on the specific prognostic stratification inferred with CELTYC. We reasoned that epigenetic alterations may have complex downstream effects on gene-expression and hence that performing GSEA ^55,56^ on genes ranked by differential expression between CELTYC’s poor and good outcome clusters would be more fruitful. This revealed enrichment of pathways well-known to impart a poor outcome, including EMT, TGF-beta signaling and angiogenesis (**Fig. 5b**). Differentially expressed genes (DEGs) between the poor and good outcome clusters also contained a significantly higher number of DMCTs than what would have been expected by random chance (**SI fig.S11a**). To see which DMCTs within DEGs were associated with the gene’s mRNA expression level we performed correlation analysis over the tumors, revealing a significant number of correlated and anti-correlated DMCT-DEG pairs (**SI fig.S11b**), with the corresponding DMCTs mapping significantly more often to either gene-body or to within 200bp upstream of the gene’s transcription start site (**SI fig.S11c**). Gene over-representation analysis using DAVID ^57^ revealed marginal enrichment of biological terms related to membrane trafficking, cell-cycle, mesothelioma, pluripotency, hepatocellular carcinoma and WNT-signaling (**SI fig.S11d, SI table S10**). It is striking that among 4 enriched genes (*FZD1, LRP5, AKT2, AXIN1*) implicated in both mesothelioma and stemness (**SI fig.S11e**), three of these (*FZD1, LRP5, AXIN1*) are key members of the canonical WNT-signaling ^58–60^ pathway, one of the key pathways whose activation has been associated with increased stemness and aggressive liver cancer ^58^. Moreover, *AKT2* has been shown to regulate WNT-signaling ^61–64^. Joint heatmaps of DNAm and mRNA expression confirmed upregulation of these genes in the poor outcome cluster, with corresponding DMCTs also displaying differential DNAm (**SI fig.S11e**). Several of these genes’ mRNA expression levels were also associated with clinical outcome when assessed individually with Cox-regressions (**SI table S11**). Of note, downstream WNT signaling pathway members *CTNNB1* and *AXIN1* were more frequently mutated in the good outcome cluster (*CTNNB1* Pclust:0.23, Gclust:0.34, *AXIN1*: Pclust:0.07, Gclust:0.08), although differences were not statistically significant. Likewise, *CTNNB1* and *AXIN1* did not display significantly different deletion/loss frequency (*CTNNB1* Pclust:0.02, Gclust:0.04, AXIN1 Pclust: 0.13, Gclust: 0.13) or amplification/gain frequency (*AXIN1* Pclust:0.33 Gclust:0.24. *CTNNB1* Pclust:0.36 Gclust:0.41), pointing towards the poor outcome phenotype being driven in part by epigenetic dysregulation of WNT-signaling.

**Figure 5:**
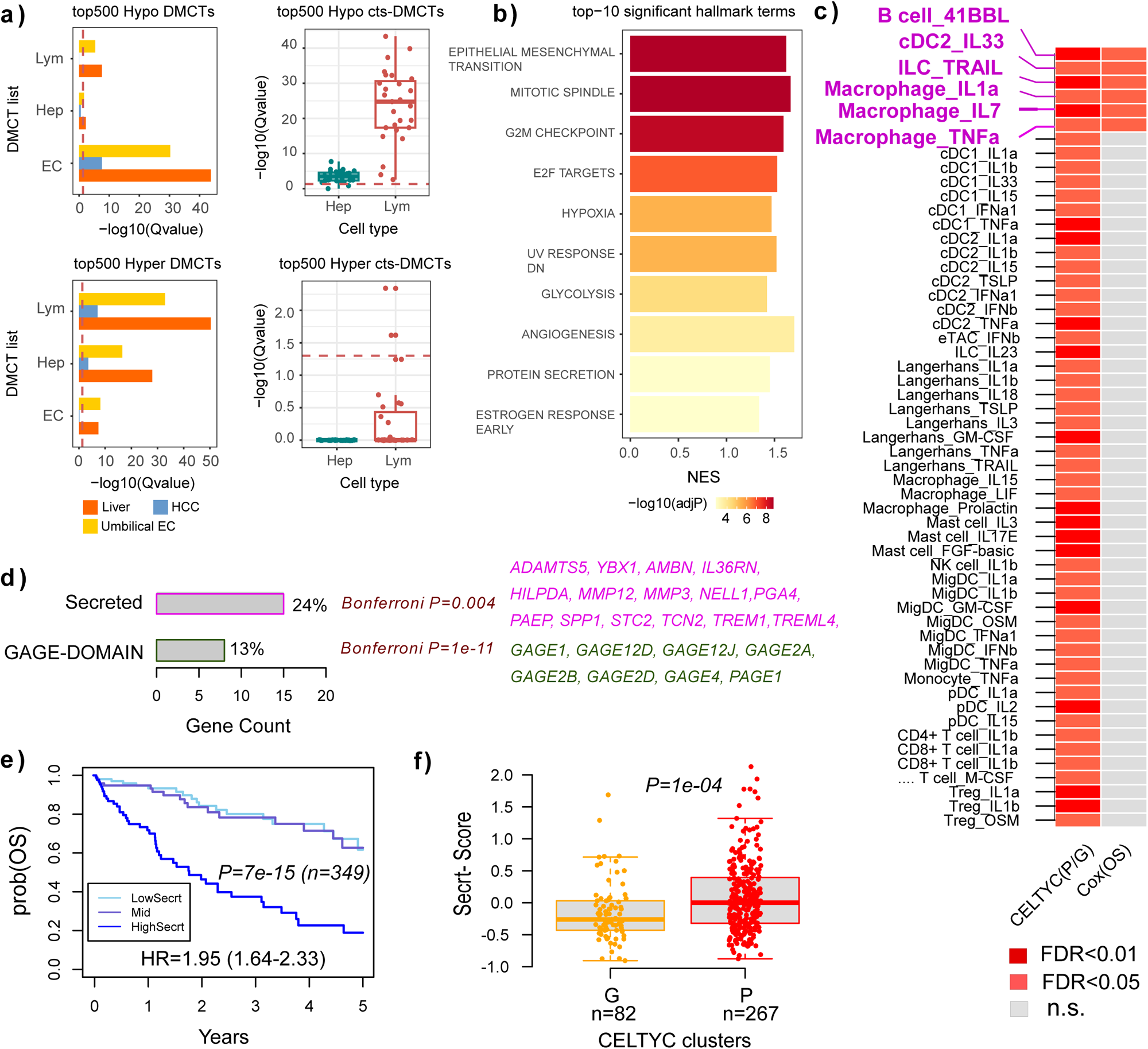
Biological interpretation of CELTYC LIHC prognostic model. **a)** eFORGE enrichment analysis results for hypo and hyper EC, Hep and Lym-DMCTs in LIHC. Left top: Barplots display significance of enrichment (-log10[Q-value]) of top-500 hypo EC, Hep and Lym-DMCTs for active transcription start sites (TSS), as defined in 3 different cell/tissue types (liver, hepatocellular carcinoma, umbilical endothelial cell); Right top: Boxplot comparing the statistical enrichment levels of the top-500 hypo Hep and Lym specific DMCTs (i.e. not shared by different cell types) in active TSS of immune cells; Left bottom: the same as left top, but for top-500 hyper DMCTs. Right bottom: the same as right top, but for top-500 hyper DMCTs. **b)** Barplot displaying the normalized enrichment scores (NES) from GSEA and significance levels (-log10[Adjusted P-value]) of genes ranked by differential expression between Pclust and Gclust for the top enriched MSigDB hallmark gene sets. **c)** Heatmap of FDR (Benjamini-Hochberg) associations of 55 cytokine signatures with poor/good (P/G) outcome CELTYC clusters and with overall survival (Cox(OS)). In all cases, associations mean higher cytokine activity in P cluster or in poor outcome. P-values were estimated from a linear model (CELTYC clusters) or a Cox-regression (for outcome). Six cytokine signatures significant in both are highlighted in magenta. Cytokine signatures are labeled by cell-type and cytokine applied to that cell-type. **d)** DAVID enrichment analysis of genes upregulated in poor outcome LIHC samples, as determined by Cox-regressions, listing the top two enriched categories. Percentage values means the percentage of genes in that term that were among the outcome associated upregulated genes. Bonferroni adjusted P-values are given alongside the enriched genes in each category. **e)** Kaplan-Meier overall survival curve for LIHC samples stratified according to lower, middle and higher tertiles of the secretory-score, computed as the average gene-expression of the 15 enriched secretory genes. Hazard ratio, 95% confidence interval and score-test P-value are given. **f)** Boxplot comparing the same secretory-score in e) between the two CELTYC clusters. P=poor outcome, G=good outcome. P-value derives from a one-tailed Wilcoxon-test.

### A cytokine secretion signature is associated with CELTYC’s poor outcome cluster

Activated WNT-signaling has been linked to increased stemness and immune evasion, promoting a more aggressive hepatocellular carcinoma phenotype ^65,66^. To explore in more depth the relevance of the immune-system component, and given the growing importance of epigenetically associated immune modulation in the tumor microenvironment ^16,67–69^, we calculated cytokine-activity scores for cytokine signatures from the “Immune Dictionary” ^70^, a large compendium of 938 immune cell-type specific cytokine-response expression signatures derived from single-cell RNA-Seq data, encompassing 17 immune-cell types and 86 cytokines (**Methods**). This revealed 55 cytokine signatures (adjusted linear model P < 0.05) correlating with the CELTYC clusters, with all 55 displaying increased activity in the poor outcome cluster. Of these 55 signatures, a total of 6 (ILC_TRAIL, cDC2_IL33, Macrophage_IL1a/IL7/TNFa and B-cell_41BBL) were also significantly correlated with overall survival (adjusted Cox-regression P < 0.05) (**Fig. 5c**). Of note, conventional type-2 dendritic cells (cDC2) are thought to be key regulators of inflammation and IL33 has generally been associated with activation of poor outcome type-2 immune responses (e.g. T-helper-2) ^71^. A signature measuring TRAIL (TNFSF10) stimulation in innate lymphocyte-cells (ILCs), a subset of highly interactive tissue-resident lymphoid cells that regulate chronic inflammation and tissue-homeostasis ^72,73^, was also associated with poor outcome. Resistance of liver cancer cells to apoptosis by TNFSF10, with TNFSF10 also promoting metastasis, has recently been documented ^74,75^, which could partly explain the association with poor outcome found here. Moreover, TNFSF10 has been shown to induce a cancer cytokine secretome ^75^, and in line with this, gene set overrepresentation analysis with DAVID ^57^ revealed a strong enrichment for a secretory phenotype among genes negatively associated with clinical outcome (**Fig. 5d**). Average expression of these secretory factors was strongly associated with clinical outcome and higher in the poor outcome CELTYC cluster (**Fig. 5e-f**). Using CIBERSORTx ^76^ to identify cell-type specific differential expression ^77^, in this instance, lymphocyte specific DEGs, we were able to confirm enrichment of secretory and T-helper-2 pathways (**SI fig.S12**). In summary, these data show that CELTYC’s prognostic clusters may in part be driven by a complex cancer cytokine secretome that promotes a type-2 immune response environment.

### CELTYC reveals cell-type specific prognostic subtypes in kidney cancer

To demonstrate that CELTYC can reveal novel subtypes in other cancer-types, we considered the case of kidney renal carcinoma (KIRC), a cancer type for which a large number of normal-adjacent and cancer samples were profiled as part of the TCGA (n=160 normal-adjacent + 319 cancers) ^78^. To estimate cell-type fractions in the KIRC samples we applied our validated HEpiDISH DNAm reference matrix, defined over a generic epithelial, fibroblast and immune cell-type ^12^ (**SI fig.S13a-b**). To independently check that these fractions are reasonable, we compared them to those obtained using a separate DNAm reference matrix built from a recent WGBS DNAm-atlas ^79^ (**Methods**), which resulted in an excellent agreement for the shared epithelial and immune cell components (**SI fig.S13c**). Applying CellDMC with the estimated cell-type fractions, we inferred epithelial, stromal (fibroblast) and immune-cell specific DMCTs (**SI table S12**), with the overwhelming majority of changes occurring in the epithelial compartment (**Fig. 6a**). We verified that clustering over these DMCTs resulted in segregation of samples by normal-cancer status (**SI fig.S13d**). Next, we clustered the cancer samples only, doing so separately over the epithelial, fibroblast and immune-cell specific DMCTs. In the case of epithelial-DMCTs, this revealed 4 optimal clusters that showed some correlation with the clusters obtained over fibroblast and immune-cell DMCTs, but which were however also clearly distinct (**Fig. 6b**). Effect sizes between clusters, as defined in the original unscaled basis, were typically in the range of 1-30% DNAm change for all DMCT defined clusters (**SI fig.S14**). Kaplan Meier analysis revealed that for all cell-types, clusters differed strongly with respect to overall survival (**Fig. 6c**). To explore this further, we collapsed the clusters for each cell-type into 3 groups of good, intermediate and poor outcome (**Fig. 6d**), based on their original KM-curve distributions (**Fig. 6c**). Although age, stage, grade and residual tumor were each strongly associated with poor outcome (**SI table S13**), for the fibroblast and immune-cell derived CELTYC-clusters the associations with overall survival remained significant upon adjustment for all of these factors (**SI table S13**). Importantly, a prognostic model built from all 3 cell-type specific clusterings significantly outperformed one based on clusters derived from ordinary differentially methylated cytosines (DMCs) (Likelihood ratio test, 2 dof, P<6e-6). This demonstrates that a reductionist cell-type specific approach can improve prognostic stratification of KIRC compared to cell-type agnostic clustering.

**Figure 6:**
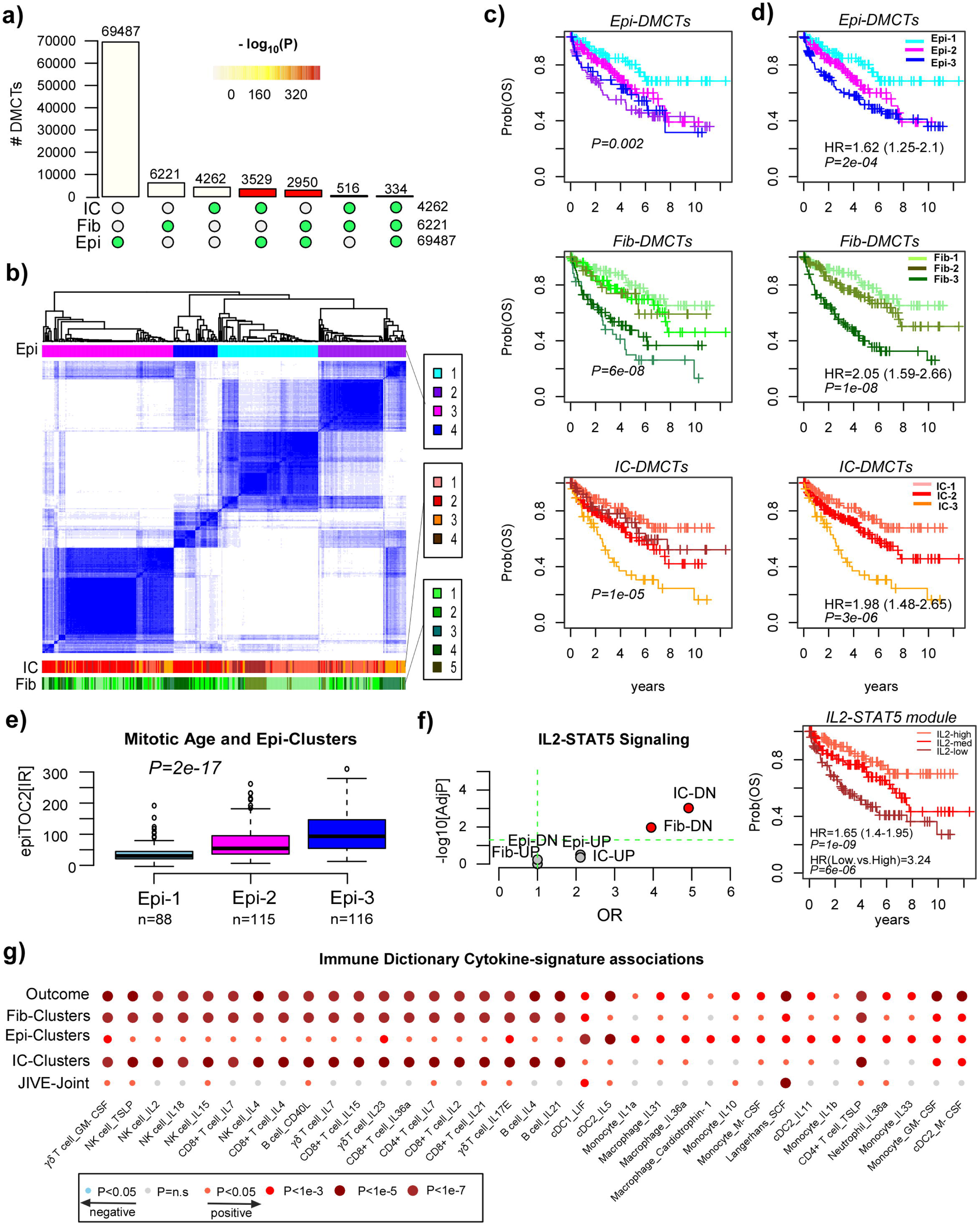
CELTYC identifies cell-type specific prognostic subtypes in KIRC. a) An upset plot displaying the number of cancer-associated DMCTs inferred in each of 3 broad cell-types (Epi=epithelial, Fib=fibroblast/stromal, IC=immune-cell) by applying the full CellDMC model to the TCGA KIRC DNAm dataset. **b)** Consensus clustering result for the optimal 4 clusters obtained by clustering over the epithelial-specific DMCTs. The distribution of corresponding immune-cell DMCT and fibroblast-DMCT derived clusters is shown in the barplots at the bottom. **c)** Kaplan Meier Analysis depicting the overall survival distributions of the corresponding clusters in b). The P-values are from a log-rank test. **d)** As c) but now stratifying the clusters into groups of poor, intermediate and good outcome according to the survival distributions shown in b). Hazard Ratio, 95% confidence interval and Chi-Square score test P-value is given. **e)** Boxplots displaying the correlation of mitotic age (as estimated using epiTOC2’s annual intrinsic rate of cell division (IR)) with the epithelial-DMCT derived clusters of d). P-value is two-tailed from a linear regression. **f) Left:** Enrichment overrepresentation analysis result for the IL2-STAT5 hallmark gene set for 6 categories of genes: genes with mRNA expression correlating positively (UP) or negatively (DN) with the three sets of poor outcome clusters in d), one for each cell-type. Scatterplot displays the Odds Ratio (OR, x-axis) and statistical significance (-log10[adjusted P-value], y-axis) from a one-tailed Fisher’s exact test. **Right:** Kaplan-Meier curves for KIRC samples stratified into 3 tertiles according to the average expression of 11 enriched IL2-STAT5 genes. Hazard Ratio (HR), 95% confidence interval and Chi-Square P-value is given from a Cox-regression of overall survival against the negative average expression of the IL2-STAT5 genes. We also display the HR and corresponding Chi-Square P-value comparing the two extreme tertiles. **g)** Balloon plot of associations between immune-cell type specific cytokine activity scores and the CELTYC-derived fibroblast, epithelial and immune-cell clusters, the clusters derived from JIVE’s joint variation matrix, as well as overall survival. In the case of the clusters, the two-tailed P-values derive from a multivariate linear regression of cytokine signature activity against ordinal cluster-number adjusting for epithelial, fibroblast and total immune-cell fractions. In the case of clinical outcome, the P-value derives from the Chi-Square score-test of a proportional hazards Cox-regression. Cytokine-signature activity was defined as the average expression of genes specified as upregulated in the signature. Each signature is labeled by cell-type and cytokine.

To study the biological significance of these KIRC subtypes we first performed an enrichment overrepresentation analysis ^80^ of genes containing cell-type specific DMCTs that are also significantly up or downregulated in cancer compared to normal-adjacent tissue (**Methods**). Testing enrichment against the highly curated cancer-hallmark set from MSigDB ^55,56^ revealed in the case of epi-DMCT genes upregulated in cancer, a strong enrichment for bivalent genes, inflammatory response, glycolysis, hypoxia, activated KRAS-signaling and EMT (**SI fig.S15a**). In contrast, downregulated genes were only enriched for bivalent and PRC2 genes, and deactivated KRAS-signaling (**SI fig.S15a**). The differentially expressed fibroblast-DMCT genes displayed less enrichment, but higher enrichment of the complement pathway, whilst the immune component displayed no enrichment due to the lower number of DMCTs in this compartment (**SI fig.S15a**). The observation that downregulated genes with underlying differential DNAm changes in the epithelial compartment are enriched for bivalent and PRC2-targets, points towards cell-proliferation as the underlying mechanism since PRC2-targets are prone to promoter DNA hypermethylation changes following cell-division ^81,82^. The concomitant enrichment for KRAS-signaling and glycolysis further points to specific oncogenic sources of increased proliferation. To explore this further, we reasoned that the CELTYC epithelial-DMCT clusters may be related to the mitotic age of the tissue ^81,83^. Estimating the mitotic age of all samples using epiTOC2, a DNAm-based mitotic clock ^83^, showed an increased mitotic age in KIRC compared to age-matched normal-adjacent tissue **(SI fig.S15b**), and confirming our hypothesis, mitotic age displayed a strong correlation with the 3 CELTYC Epi-clusters and clinical outcome (**Fig. 6e**). Interestingly, whilst the CELTYC Fib-clusters also displayed a linear correlation with mitotic age, the pattern was non-linear for the IC-clusters, with an increased mitotic age only evident for the poorest outcome IC-cluster (**SI fig.S15c**). Since prognostic separability was highest in the fibroblast and immune-cell compartments (**Fig. 6d**) and in order to gain more power in our enrichment analysis, we performed differential expression analysis by identifying genes whose expression correlates most strongly with the corresponding CELTYC clusters graded by clinical outcome (**Methods**). Whilst this revealed similar enrichment patterns for the Epi-clusters, a striking observation was the enrichment of IL2-STAT5 signaling, specifically for genes downregulated in the poor outcome IC-clusters (**Fig. 6f**). This enrichment was driven by 11 genes that are activated by STAT5 upon IL2-stimulation (*BCL2, CDC42SE2, GBP4, IRF6, ITGA6, LRRC8C, PLPP1, PRKCH, RHOB, SHE, SWAP70*) ^56^. We verified that all 11 displayed significant associations with overall survival (**SI table S14**), with low average expression conferring poor outcome (HR=1.65 with 95%CI: 1.40-1.95, P=10^-9^, **Fig. 6f**). We also computed cytokine activity scores for 333 signatures from the Immune Dictionary compendium ^70^ that had sufficient gene-representation in the TCGA data (**Methods**), observing how associations were strongest for the CELTYC immune-cell and fibroblast clusters (**SI fig.S16a, SI table S15, Fig. 6g**). Almost all associations were positive, i.e. high cytokine-activity correlates with poor outcome clusters and overall survival (**Fig. 6g, SI fig.S16a**). Interestingly, whilst the strongest cytokine associations with the CELTYC immune cell clusters generally mapped to T-cell lymphocyte subtypes (e.g. IL2 on CD8+ T-cells), the opposite was true for the associations with the epithelial clusters which were dominated by myeloid cells (e.g. dendritic cells, macrophages) (**Fig. 6g, SI fig.S16b**). We verified that all three cell-type specific prognostic models were independent of recent single-cell RNA-Seq derived prognostic macrophage signatures ^84^ (**SI fig.S17**, **Methods**). That increased IL2 activity in CD8+ T-cells is associated with poor outcome (**Fig. 6g**) could be consistent with recent reports that IL2 acts to induce CD8+ T cell exhaustion within tumor microenvironments ^85^ and that CD8+ T-cell exhaustion is a key factor underlying metastasis and poor outcome ^26^. Overall, these data indicate how CELTYC can dissect poor clinical outcome into separate epithelial and immune-cell components, reflecting increased mitotic age and altered cytokine signaling, respectively. Of note, by applying JIVE to extract joint and cell-type specific variation matrices, we observed that inferred consensus clusters displayed greatest prognostic separability for the joint variation (**SI fig.S18**). This suggests that, although we have identified distinct cell-type specific prognostic subtypes, there are complex coordinated functional epigenetic changes between cellular compartments.

### Prognostic synergy and validation in KIRC

Since CELTYC has been able to dissect mitotic age and IL2-STAT5 signaling, both associated with poor outcome, into their underlying cellular compartments, we next asked if these two processes may synergize to yield stronger prognostic models. To this end, we performed combinational clustering, i.e. we stratified all KIRC samples into 9 groups based on the CELTYC epithelial (Epi1,Epi2,Epi3) and immune-cell (IC1,IC2,IC3) clusters and generated KM-curves for each (**Fig. 7a**). This revealed strong synergy, in the sense that those samples with highest mitotic-age (Epi3) and lowest IL2-STAT5 signaling (IC3) had the worst clinical outcome, whereas samples with lowest mitotic age and highest IL2-STAT5 signaling (Epi1-IC1) displayed the best outcome. Stratifying all KIRC samples into these two subgroups and the rest, revealed an approximately 70% difference in overall survival 10 years after diagnosis (**Fig. 7b**). The Odds Ratio of a death event in the Epi3-IC3 group compared to Epi1-IC1 was 68 (Fisher test P=4e-10). To formally test for prognostic synergy we performed likelihood-ratio tests comparing the prognostic model defined by combinatorial clustering (**Fig. 7a**) to the ones defined separately by the epithelial and immune-cell clusters (**Methods**), revealing that the combinatorial model significantly improved prognostic stratification (**Fig. 7c**). Prognostic synergy was also seen when comparing the combinatorial ordinal clusters (**Fig. 7b**) to the ordinal epithelial and immune cell clusters (**Fig. 7c, Methods**).

**Figure 7:**
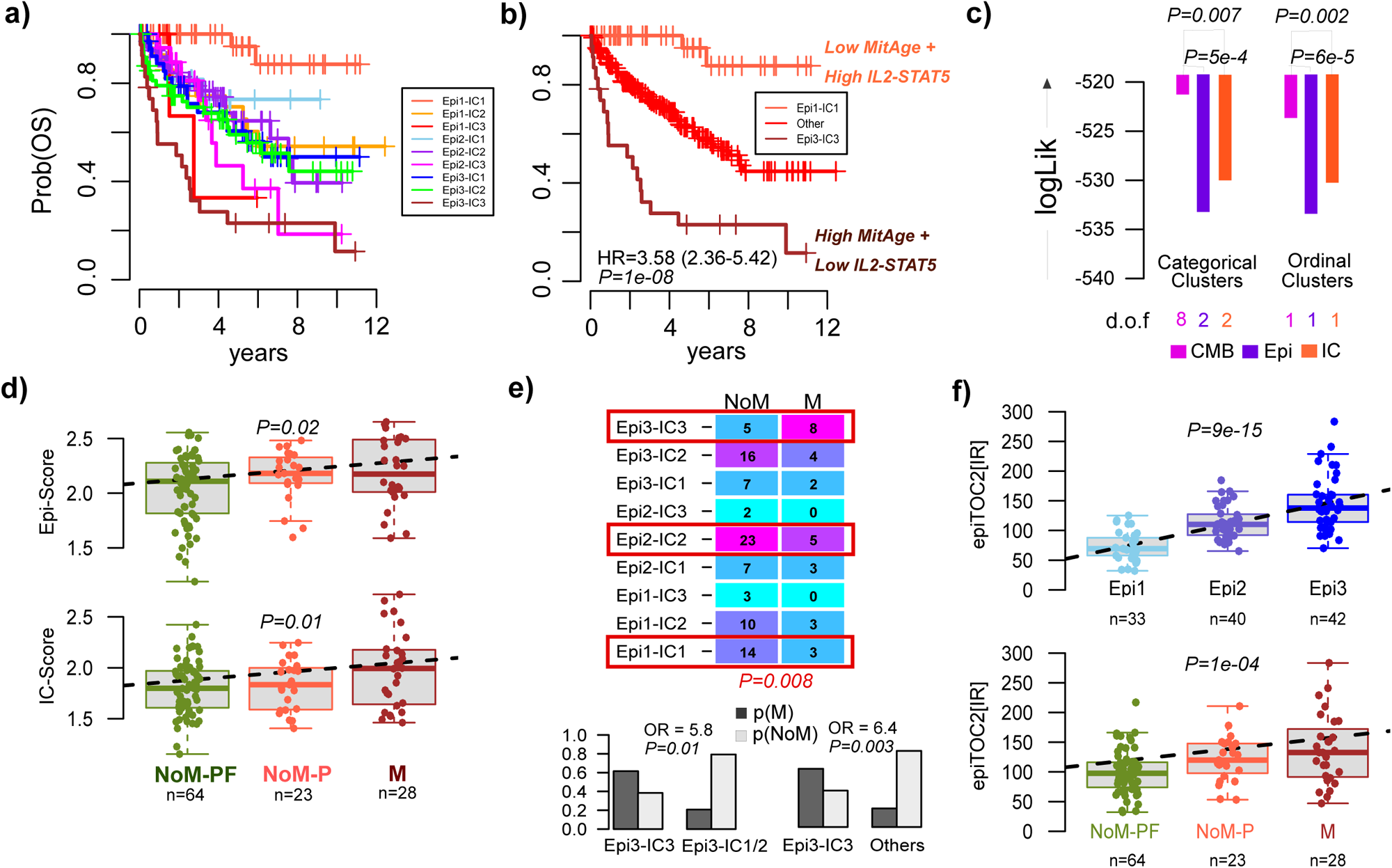
Combinatorial indexing of CELTYC clusters reveals prognostic synergy. **a)** Kaplan Meier overall survival plots of the 9 subgroups defined by combinatorial indexing of the 3 CELTYC epithelial and 3 CELTYC immune-cell clusters. **b)** Same as a) but grouping samples into 3 groups with “Other” labeling all subgroups other than Epi3-IC3 and Epi1-IC1. Hazard Ratio (HR), 95% confidence interval and Chi-square score test P-value from a Cox regression of overall survival against ordinal cluster number (1=Epi1-IC1, 2=Other, 3=Epi3-IC3) is given. **c)** Barplots depict the log-likelihoods of Cox-regression models for overall survival where the covariate is either categorical or ordinal. For the categorical case, the covariates are the combinatorial indexing of epithelial and immune cell CELTYC clusters as defined in a) (CMB), the categorical epithelial CELTYC clusters (Epi) and the categorical immune-cell CELTYC clusters (IC). For the ordinal case, the covariates are the 3 ordinal combinatorial clusters as defined in b) (CMB), the 3 epithelial CELTYC clusters treated as ordinal (Epi) and the 3 immune-cell clusters also treated as ordinal (IC). The number of degrees of freedom (dof) is given for each model. For the categorical case, the P-values are derived from a one-tailed ChiSquare Likelihood Ratio test with dof given by the difference between the two models being compared. For the ordinal case, the P-values represent the relative probability of the IC or Epi model being better than the CMB-model. **d)** Validation of the Epi and IC CELTYC cluster-predictors in an independent ccRCC DNAm dataset. NoM-PF: non-metastatic progression-free, NoM-P: non-metastatic and progression, M=metastatic. P-value is from a linear regression of the predictors scores against disease stage. **e)** Distribution of non-metastatic and metastatic events among the 9 groups obtained by combinatorial indexing of the predicted Epi and IC-clusters in the independent dataset. Barplots compare the probabilities of a metastatic event among different subgroups, with the Odds Ratio and one-tailed P-value derived from a Fisher’s exact test on sample numbers. **f)** Boxplots of the epiTOC2 mitotic-age score (average lifetime intrinsic rate of stem-cell division which is naturally age-adjusted) against predicted Epi-cluster and disease stage in the independent dataset. P-value is from a linear regression treating Epi-cluster and disease stage as ordinal variables.

To test whether this prognostic synergy is also seen in independent cohorts, we first applied a 5-fold cross-validation strategy to build linear Elastic Net ^86^ DNAm-predictors for the epithelial and immune-cell CELTYC clusters, using only the corresponding cell-type specific DMCTs as input in the training process (**SI table S16-17, SI fig.S19a, Methods**). This resulted in two separate predictors for assigning samples to one of the 3 immune-cell or one of the 3 epithelial-cell CELTYC clusters (the IC-predictor and Epi-predictor). As required, the scores from these two predictors were correlated with clinical outcome in the TCGA KIRC samples (**SI table S18**). To validate the predictors, we applied them to an independent Illumina 450k dataset ^87^ of 132 clear-cell renal cell carcinomas (ccRCC) and 12 controls (**Methods**). Although exact overall survival information was not available for this cohort, samples were stratified into 3 separate categories (non-metastatic and progression-free, non-metastatic that progressed and metastatic at diagnosis), allowing for validation. Both the IC as well as the Epi-predictors yielded scores that increased with disease progression (**Fig. 7d**). Stratifying the ccRCC patients into nine separate prognostic groups according to their predicted IC and Epi-scores (**Methods**), revealed a significant differential distribution of metastatic events, specially when comparing the predicted poor outcome Epi3-IC3 cluster to the rest (**Fig. 7e**). Of note, none of the other Epi3 or IC3 clusters revealed a preponderance of metastatic events (**Fig. 7e**), supporting the view that it is simultaneous high mitotic age and low IL2-STAT5 signaling that drives poor outcome. In support of this, mitotic age of the ccRCC samples displayed a very strong correlation with the Epi-predictor and clinical outcome (**Fig. 7f**), and average DNAm of promoter CpGs mapping to IL2-STAT5 genes was also increased in the IC-3 cluster, albeit only marginally so (**SI fig.S19b**). Overall, these data validate the synergistic poor-outcome effect of high mitotic-age and low IL2-STAT5 signaling in KIRC.

### CELTYC is applicable to large RRBS datasets

In the previous applications to LIHC and KIRC, the DNAm data had been generated with Illumina 450k beadarrays, which summarize DNAm at the level of individual CpGs. To demonstrate that CELTYC can be applied to other technologies, we considered a large reduced representation bisulfite sequencing (RRBS) breast cancer dataset ^88^ encompassing 1479 breast cancers and 231 normal-adjacent tissue specimens, and where the DNAm data has been summarized at the gene-promoter level (**Methods**). As proof-of-concept, we considered the case of luminal estrogen receptor positive (ER+) breast cancer, as this is the predominant subtype with 797 samples. We used our EpiSCORE DNAm reference matrix for breast tissue, encompassing representative DNAm profiles for basal, luminal, adipocytes, endothelial, fibrobast, lymphocyte and macrophage cells, estimating corresponding cell-type fractions in all 1028 (231+797) samples (**SI fig.S20a**). CellDMC predicted most cancer-associated cell-type specific differential methylation (**SI table S19**) to occur in the luminal compartment, followed by changes in endothelial cells and lymphocytes (**SI fig.S20b**). DNAm changes in promoters displayed clear preferential hypermethylation in the luminal and lymphocyte compartments but preferential hypomethylation in endothelial cells (**SI fig.S20b**). Because differentially methylated gene promoters (DMGs) largely overlapped between cellular compartments, for interpretability we next restricted to cell-type specific DMGs, performing separate clustering over each cell-type specific set of DMGs. Clustering over the luminal-specific DMGs led to an optimal 3-cluster solution that was prognostic, with two clusters mapping broadly to the luminal A and B subtypes, and with the third cluster displaying an intermediate outcome phenotype (**SI fig.S20c-d**). Clustering over the endothelial cell and lymphocyte DMGs also led to prognostic models, but not outperforming the transcriptomic based (lumA vs lumB) prognostic model. Although combinatorial clustering also did not reveal any improved prognostic models, we aimed to validate the cell-type specific DMGs. In the case of the hypermethylated gene promoters occurring in the luminal compartment, these were once again strongly enriched for PRC2-marked and bivalent genes (**SI fig.S20e**), similar to the pattern observed in the epithelial compartment of KIRC. We reasoned that the CELTYC luminal derived clusters could thus reflect differences in cell proliferation and mitotic age, which was validated by application of epiTOC2 (**SI fig.S20d**), further confirming that a significant portion of the DNAm landscape in breast cancer is driven by cell-proliferation ^88,89^. In contrast to the luminal compartment, the strongest enriched terms for lymphocyte-DMGs were enriched for immune cell signatures, including interferon alpha and gamma responses and CD4T-cells, whilst for the endothelial compartment, we observed marginal enrichment for genes expressed in fetal endothelial cells ^90^ (**SI fig.S20d**). Thus, although in this instance, CELTYC did not lead to prognostic models that outperform established ones, it does clearly identify prognostic signatures within individual cellular compartments, thus demonstrating that it is applicable to other cancer-types and technologies.

## Discussion

Here we have advanced the concept of cell-type specific combinatorial clustering (CELTYC), demonstrating, in two different cancer-types, how it can refine and improve prognostic cancer classifications. Conceptually, that such improvements should be possible is highly plausible, since current cancer classifications are generally speaking derived from bulk-tissues that are composed of many different cell-types, with the substantial inter-subject variations in tissue-composition overly confounding or masking cell-type specific prognostic associations. Whilst cellular composition of tissues is also highly informative of disease subtypes and prognosis, the reductionist, cell-type specific, CELTYC approach allows construction of improved prognostic models by combinatorial indexing of the cell-type specific clusters, as shown here for KIRC. This in turn can also help elucidate the biological meaning of cancer subtypes. For instance, in the case of KIRC, CELTYC was able to correctly dissect and assign two distinct biological processes associated with poor clinical outcome (increased mitotic-age/cell proliferation and reduced IL2-STAT5 signaling), to the corresponding epithelial and immune-cell compartments, respectively. This allowed construction of a significantly improved prognostic model based on combinatorial indexing of the epithelial and immune-cell clusters, which is not attainable with standard methods that do not use cell-type deconvolution. Although we were able to validate the synergistic effect of high mitotic age and low IL2-STAT5 signaling in an independent ccRCC cohort, by no means are we arguing that there are no other immune-signaling axes contributing to poor outcome in KIRC. Indeed, application of the cytokine Immune Dictionary revealed many cytokine lymphocyte signatures contributing to poor outcome in KIRC, including e.g. IL2 stimulation of NK and CD8+ T-cells, or IL-7 stimulation of CD4+, CD8+ and γδ T-cells. Although IL2-signalling is normally associated with favorable outcome, IL2 can also amplify T regulatory cells ^91,92^ and a recent study has implicated IL2 in stimulating CD8+ T-cell exhaustion ^85^, with CD8+ T-cell exhaustion emerging as the key marker of poor outcome in KIRC/ccRCC ^26^. That low IL2-STAT5 and high IL2 T-cell signaling are simultaneously associated with poor outcome KIRC samples may indicate a complex intricate rewiring of IL2-signaling, or alternatively, that the two IL2 signature states may not necessarily be operative in exactly the same poor outcome samples. Another interesting finding was the differential association of cytokine signatures across the immune and epithelial-cell compartments, with myeloid signatures almost exclusively associated with the mitotic-age clusters defined by epithelial-DMCTs. This may indicate an intricate paracrine signaling between poor outcome IL1, IL10 and IL36a macrophage polarization ^93–96^ and the proliferation-state of tumor cells, as demonstrated recently in the case of colon cancer ^97^.

The application of CELTYC to LIHC also led to an improved prognostic model, which was validated at the mRNA level in 2 independent LIHC datasets. Although the association with poor outcome was driven by lymphocyte-DMCTs, many overlapped with respective DNAm changes in the epithelial and endothelial cell compartments, suggesting highly consistent and coordinated epigenetic changes across cell-types. Consistent with this, differentially expressed genes correlated with DMCTs were enriched for terms normally associated with a poor outcome including stemness and activated WNT-signaling. The latter enrichment was particularly striking, involving not only the co-receptors *FZD1* and *LRP5*, but also an element (*AXIN1*) of the beta-catenin destruction complex, as well as external regulators of WNT-signaling such as *AKT2* ^61–64^. Whilst the role of WNT-signaling in hepatocellular carcinogenesis is well-known ^58–60^, with *CTNNB1* and *AXIN1* frequently mutated or amplified, it is worth stressing that these specific genomic alterations did not display variable frequencies between the poor and good outcome clusters, suggesting that other mechanisms are contributing to differential prognosis. In line with this, our work highlights the potential importance of differential DNAm in the gene-body or promoter of genes like *FZD1, LRP5, AXIN1* and *AKT2* in contributing to the differential prognosis. We note that the identification of *FZD1* and *LRP5* is particular interesting given earlier work demonstrating how epigenetic dysregulation of genes in cancer happens preferentially in the extracellular and membrane receptor domains, in contract to genetic dysregulation which preferentially targets intracellular domains ^67^. Altered WNT-signaling has also been linked to an altered immune tumor microenvironment and immune evasion ^65,66^. Cytokine signature analysis revealed the presence of 6 signatures that could potentially contribute to the poor outcome phenotype, alongside other well-known processes such as EMT and angiogenesis. Among these signatures, it is worth highlighting again the TNFSF10-CD8+ T-cell stimulation signature, as targeted TRAIL (TNFSF10) therapy is being extensively explored in clinical trials ^98^. Intriguingly, TRAIL has been associated with a tumor promoting secretory phenotype ^75^, and consistent with this GSEA revealed a highly significant enrichment of secretory proteins including matrix metalloproteinases. Interestingly, the same GSEA revealed an even stronger enrichment for genes with GAGE domains (cancer and testis specific antigens), which have been implicated with metastasis and poor outcome in a range of different cancer types and are potential targets for immunotherapy ^99,100^.

It is clear that CELTYC is particularly useful (and necessary) in the context of DNAm data. For mRNA expression, future single-cell approaches on large numbers of clinical samples can more directly lead to cell-type specific disease subtyping, as shown recently in the context of eQTLs ^101^. For DNAm data however, this is not going to be feasible in the foreseeable future. Thus, critical to help realize the potential and value of CELTYC in the DNAm context is the development of highly accurate tissue-specific DNAm reference matrices at high cellular resolution as well as sufficiently large DNAm datasets to ensure that algorithms such as CellDMC can capture most of the DMCTs in each underlying cell-type. As shown here, EpiSCORE’s DNAm-atlas or the more coarse-grained HEpiDISH DNAm reference matrix, can help dissect DNAm patterns into the DNAm changes displayed by different cell-types, but the cellular resolution is still limited to a few cell-types. This is partly because these DNAm reference matrices need further improvement, but also because inference of DMCTs over more cell-types requires concomitant increases in sample size. As demonstrated here, one would require at least 400 samples, ideally with balanced numbers of normal and disease samples, to have enough sensitivity to detect a reasonable number of true-DMCTs in at least one or two of the different cell-types in the tissue. Thus, larger DNAm datasets will be required to more reliably identify a larger set of DMCTs in three or more cell-types. In line with this, GSEA did not always reveal clear segregated enrichment of cell-type specific biological terms among the respective cell-type specific DMCTs, highlighting that sensitivities are relatively low and FDR’s relatively high. On the other hand, the availability of matched RNA-Seq data can help sharpen GSEA results, rendering cell-type specific enrichments, as illustrated by the enrichment of bivalent PRC2-target and IL2-STAT5 signaling genes in the epithelial and immune-cell compartments of KIRC, respectively.

Although our focus has been on TCGA DNAm data that was generated with Illumina 450k beadarrays and which was analyzed with CELTYC at the level of individual CpGs, it is worth pointing out that CELTYC is straightforwardly applicable to other DNAm technologies and for DNAm data summarized at the level of regulatory regions, as illustrated here for RRBS data and gene promoters. This demonstration was also performed in the context of a different cancer-type (breast cancer). In future, it might be interesting to extend CellDMC and CELTYC to infer cell-type specific differentially methylation regions, which could lead to improved robustness over the inferences performed at the individual CpG-level.

In summary, we have demonstrated that cell-type specific combinatorial clustering of DNAm data can lead to distinct and improved prognostic models in cancer, shedding new biological insights and formulating new hypotheses regarding the molecular pathways driving these models. As such, we envisage that CELTYC will be of great value to uncover clinically relevant subtypes in other cancer-types where cell-type heterogeneity would otherwise mask them.

## Methods

### Simulation models

To provide a rationale for the CELTYC procedure, we devised two separate simulation models. In one model, we considered mixtures of 3 sorted immune-cell subtypes (139 Neutrophils, 139 Monocytes and 139 CD4+ T-cells) using Illumina 450k data from BLUEPRINT ^33^. Mixture weights, i.e. the cell-type fractions, were chosen from realistic estimates by applying EpiDISH ^35^ to the 656 whole blood Illumina 450k DNAm data from Hannum et al ^102^. Because the simulated mixtures only contain 3 cell-types, estimated fractions for all lymphocytes were added together to yield the weight for the CD4+ T-cell component. Likewise, the eosinophil and neutrophil fractions were added to yield the neutrophil/granulocyte component. We generated a total of 139 mixtures, 70 representing “controls” and 69 representing “disease”. Before mixing the DNAm profiles of sorted cells together, we selected 100 random CpGs with ultra-low DNAm (beta < 0.2) across all 139 monocyte samples and high DNAm (beta > 0.8) across the other two (Neu+CD4T). For the 69 cases, we then altered DNAm of the monocyte profiles at these 100 loci by drawing them from a beta-distribution with parameters a=8, b=2, i.e. from a beta-distribution with mean 0.8. Thus, this is a scenario of a relatively big effect size where loci undergo on average >0.6 DNAm changes. For the 139 simulated mixture dataset, we then applied SVD and hierarchical clustering, as well as CellDMC ^39^ with estimated cell-type fractions using EpiDISH to infer cell-type specific DMCs (DMCTs).

The second simulation model simulates lung-tissue mixtures by mixing together sorted bronchial epithelial cells (BECs, n=108) (Magnaye et al ^42^) from 71 adult children with asthma and 37 controls, with the 139 monocytes, 139 neutrophils and 139 CD4T-cells from BLUEPRINT. The Magnaye et al raw idat files were downloaded from GEO with accession number GSE210843 and processed with R package *minfi* ^103^ to retain only probes with significant detection P values (P<0.05) across all samples. Subsequently, type-2 probe correction was performed with BMIQ^104^. Mixture weights were determined by applying EpiSCORE’s lung DNAm reference matrix ^34,37^ to the (n>200) eGTEX EPIC DNAm dataset ^43^ to infer realistic epithelial, granulocyte, monocyte and lymphocyte fractions, which were then used to mix together the BECs, neutrophils, monocytes and CD4T-cells. A total of 108 mixtures were generated with case/control status determined by the asthma-status of the original 108 BEC samples. Before mixing the sorted cells together, we defined a “ground-truth” set of 1000 asthma-DMCs (FDR<0.05) by comparing the BECs of 71 asthma-cases to the 37 controls using the limma empirical Bayes framework ^105,106^. For the 108 simulated mixtures, we then applied CellDMC, estimating cell-type fractions with HEpiDISH ^107^, a hierarchical recursive version of EpiDISH that can estimate epithelial, stromal and immune-cell subfractions for any tissue-type. We assessed the sensitivity to detect the 1000 asthma-DMCs among CellDMC’s BEC-DMCTs. Clustering was done on the standardized residual matrix over the BEC-DMCTs after regressing out the cell-type fractions. To benchmark the CELTYC performance, we also did clustering over the top 3000 most variable CpGs.

### Power calculation for in-silico mixtures

The second simulation model above provides the basis for a power calculation in order to assess the sample sizes needed for CELTYC/CellDMC to work. In order to expand the simulation model to arbitrarily large sample size, we took a parametric approach where for each CpG we learned the *(a,b)* parameters of the beta-distribution describing the DNAm values over the 37 controls. For the 1000 ground-truth asthma DMCs, the beta-distributions over the asthma cases were also learned. We also inferred the parametric beta distributions for all CpGs in the sorted immune cell-types from BLUEPRINT. Then we used the R function *rbeta* with the derived *(a,b)* parameters to construct sorted BEC, monocyte, neutrophil and CD4T-cell samples for a given number *n* of controls and an equal number *n* of asthmatic cases, where the parameters *(a,b)* differ between BEC cases and controls for the 1000 ground-truth DMCs, but not for the immune-cell types. The cell-type fractions (CTFs) used to generate the mixtures were drawn as before from the estimated CTFs of the 223 eGTEX lung samples (if the sample size of the mixtures is larger than 223, we sampled CTFs with replacement). We then considered two different CellDMC models: the full conditional model that includes interaction terms for all cell-type fractions (as described by Zheng et al. ^39^) and a marginal unconditional model where only one interaction term for one cell-type fraction (BEC) is included. These two separate models were applied to simulated mixture datasets of increasing sample size. For each sample size, the number of BEC-DMCTs (FDR<0.05) overlapping the 1000 ground truth asthma DMCs were used to calculate sensitivity and FDR. A total of 5 Monte-Carlo runs were performed at each sample size, which was sufficient as variation between runs was not substantial.

### Cell type specific clustering (CELTYC)

CELTYC aims to subtype disease samples by taking into consideration the molecular (DNAm) changes that are specific and/or joint to different cell-types in the tissue under consideration. The first step in CELTYC is to estimate the proportions of all main cell-types within a tissue. For solid tissues, we use either the EpiSCORE algorithm and its associated DNAm atlas of tissue-specific DNAm reference matrices^34^, or HEpiDISH ^12^. For blood, we use EpiDISH ^35,36^. EpiSCORE/HEpiDISH/EpiDISH were run on BMIQ-normalized DNAm data with “RPC” method and 500 iterations. In the second step, the CellDMC algorithm is used to identify CpGs with significant cell-type specific DNAm changes in relation to disease status, i.e. disease-associated DMCTs. Once DMCTs in each cell-type have been inferred, the union of all DMCTs can be partitioned into those that are common to all cell-type of interest, those that are shared between any given combination of cell-types of interest, and finally those that are unique to a given cell-type. The third step then involves clustering the disease samples only over these different DMCT-subsets, or alternatively one can use JIVE (Joint and Individual Variation Explained)^40,41^ to “cluster” over any desired number of DMCT-categories. Before clustering or JIVE, we regress out cell type proportions from the original BMIQ DNAm matrix, followed by z-score standardization of the residual matrix. This results in standardized residual matrices, one for each DMCT-subset, as defined earlier. Clustering and inference of optimal cluster number can then be performed over any desired DMCT-subset using ConsensusClusterPlus ^45^. Alternatively, any number of these residual matrices can be analyzed together using JIVE: this will extract out components of joint variation across all input residual matrices, as well as components of individual variation that are unique to each residual matrix. For instance, if we have 3 cell-types (A,B,C), we may be interested in the four residual matrices defined by the 3 DMCT-subsets unique to each cell-type (A,B,C) and the one DMCT-subset for the DMCTs common to all 3 cell-type. In general, let R_All_ denote the standardized residual matrix defined over the common/shared DMCT-subset, and let R_t_ denote the corresponding residual matrix over the DMCTs unique to cell-type *t.* JIVE then performs the following matrix decomposition:

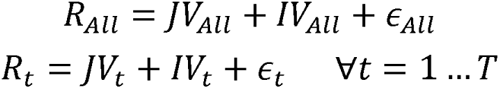

under the constraint that 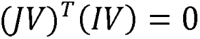 is true for each DMCT category. In practice, JIVE works in an iterative fashion, first inferring the joint variation matrix as an SVD rank *rJ* approximation obtained by stacking-up together all residual matrices, i.e. by applying SVD to 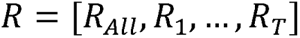, subsequently estimating the individual variation (*IV*) matrices by performing rank r_i_ SVD approximations to the residual matrices with the estimated joint variation removed. In the 2^nd^ iteration a new stacked-up matrix can be constructed by subtracting the estimated *IV* matrices from the residual matrices, subsequently reapplying SVD to this new stacked up matrix. We use the method implemented in *R.jive* ^41^ for automatic estimation of the ranks for the joint and individual variation matrices. Of note, in the example above of 3 cell-types (A,B,C), the joint variation matrix would describe variation that is common to the shared-DMCT subset and all unique DMCT-subsets, whilst the *IV*-matrices would describe variation truly unique to each DMCT-subset.

### Illumina DNAm datasets

*TCGA:* The SeSAMe ^108^ processed Illumina 450k beta value matrices were downloaded from Genomic Data Commons Data Portal (https://portal.gdc.cancer.gov/) with TCGAbiolinks ^109^. Data was downloaded and analyzed for the following cancer types: liver hepatocellular carcinoma (LIHC) and kidney clear cell renal cell carcinoma (KIRC). We did further processing of the downloaded matrices as follows: For each cancer type, probes with missing values in more than 30% samples were removed. The missing values were then imputed with impute.knn (k=5) ^110^. Type-2 probe bias was adjusted with BMIQ ^104^. Technical replicates were removed by retaining the sample with highest CpG coverage. Clinical information of TCGA samples was downloaded from Liu et al ^111^.

*Degerman:* In the case of KIRC/ccRCC, we used an independent Illumina 450k DNAm dataset of 132 ccRCC samples and 12 kidney controls ^87^. Briefly, the series data matrix file and sample annotations were downloaded from GEO (GSE113501). We only selected probes with full coverage over all samples, resulting in 391,062 probes. Provided data was already adjusted for type-2 probe bias. Of the 132 ccRCC samples, 115 had clinical outcome information available. Clinical outcome was provided as Non-metastic progression-free (n=64), non-metastatic progression (n=23) and metastatic (n=28).

*Hannum:* This is a whole blood Illumina 450k DNAm dataset of 656 samples ^102^. Data was downloaded from GEO under accession number GSE40279 and was normalized as described by us previously ^112^.

*eGTEX:* We downloaded the Illumina EPIC DNAm dataset for lung tissue from GEO under accession number GSE213478. Briefly, we downloaded the file “GSE213478_methylation_DNAm_noob_final_BMIQ_all_tissues_987.txt.gz”, which contains the already NOOB+BMIQ normalized DNAm dataset.

*BLUEPRINT:* We analyzed Illumina 450k DNAm data from BLUEPRINT ^33^, encompassing 139 monocyte, 139 CD4+ T-cell and 139 neutrophil samples from 139 subjects. This dataset was processed as described by us previously ^113^.

### Estimating cell-type fractions

In this work we estimate the proportions of all main cell-types within tissues from the TCGA using our validated EpiSCORE algorithm ^37^ and its associated DNAm atlas of tissue-specific DNAm reference matrices ^34^. This atlas comprises DNAm reference matrices for liver (5 cell-types: hepatocytes, cholangiocytes, endothelial, Kupffer, lymphocytes) and lung (7 to 9 cell-types: alveolar epithelial, basal, other epithelial, endothelial, granulocyte, lymphocyte, macrophage, monocyte and stromal). EpiSCORE was run on the BMIQ-normalized DNAm data from the TCGA with default parameters and 500 iterations. In the kidney tissue datasets (KIRC/ccRCC) we applied the HEpiDISH DNAm reference matrix, defined over a generic epithelial, fibroblast and immune-cell. This was done because the EpiSCORE kidney DNAm reference matrix was not extensively validated. We benchmarked the HEpiDISH DNAm reference matrix with another one built directly from the WGBS DNAm-atlas of Loyfer et al ^79^, encompassing 4 broad cell-types (epithelial, endothelial, fibroblast and immune-cell). Briefly, this latter DNAm reference matrix was built by identifying highly cell-type specific genes, following our previous procedure ^35^. First, we summarized DNAm values at the level of gene-promoters demanding at least 10 read coverage per CpG promoter. For the immune-cells, we had a total of 47 subtypes. For endothelial cells, we took the endothelials profiled in kidney and pancreas, yielding a total of 10 endothelial samples. For fibroblasts, we took all 7 available fibroblast samples. For the epithelial-cell component, we took the 8 available kidney podocyte samples. Subsequently, we performed limma to identify genes significantly hypomethylated in one cell-type compared to the other 3, selecting those with FDR<0.05, and ranking them by average difference in DNAm. We selected the top-50 for each cell-type, resulting in a 174 gene x 4 cell-type DNAm reference matrix.

### Other prognostic stratifications of LIHC

We used the 96 CpG DNAm signature from Ganxun Li et al ^46^ to classify TCGA LIHC samples into 3 subgroups, following the procedure of their paper. Briefly, we performed consensus clustering (*ConsensusClusterPlus*) of the DNAm profiles over the 96 CpGs, reproducing the 3-cluster solution, and marking the hypermethylated group as the predicted poor prognosis group. A separate mRNA-expression prognostic classification of LIHC was presented by Hoshida et al ^47^, which identified 3 subgroups with 2 displaying significantly poor prognosis. To reproduce the Hoshida classification, bulk mRNA expression profiles of TCGA LIHC tumors were clustered using *ConsensusClusterPlus* using the provided prognostic expression signatures for the 3 subtypes, but this resulting in an optimal 2-cluster solution with well-defined predicted poor and good outcome groups. Boyault et al. ^48^ proposed another 6 subtype classification based on expression data from 5 sets of marker genes (G1, G2, G3, G5 and G6). Here, we used ConsensusClusterPlus to cluster the TCGA LIHC samples over these genes, which resulted in an optimal 3 cluster-solution. The iCLUSTER classification annotations for TCGA LIHC samples were provided in the supplementary material (downloaded from https://ars.els-cdn.com/content/image/1-s2.0-S0092867417306396-mmc1.xlsx) from The Cancer Genome Atlas Research Network, and the Immune classification annotations for TCGA LIHC samples were provided in the supplementary material (downloaded from https://ars.els-cdn.com/content/image/1-s2.0-S1074761318301213-mmc2.xlsx) by Thorsson et al ^16^.

### Comparing different prognostic models in LIHC

To formally test that the CELTYC prognostic model defined by lymphocyte specific DMCTs (LC2 vs LC1+3) is a better prognostic model than the ones defined by other methods, we used a likelihood-based strategy that estimates relative probabilities for the respective models being true ^114^. First, we compute the Akaike Information Criterion (AIC) values for each model *m:*

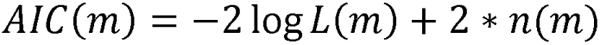

where *L(m)* is the model partial likelihood (derived from the Cox regression) and *n(m)* is the number of parameters of model *m.* Since in this case, all models have the same number of parameters, we can ignore this term. Then, for two models *k* and *j* we define

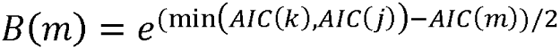

where *m* can be either *k* or *j.* Finally, the relative probability of the two models being true is given by

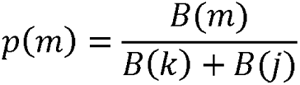

For instance, for the lymphocyte specific CELTYC model (LC), its relative probability of being true relative to the model defined by Hoshida clusters (HSD), would be:

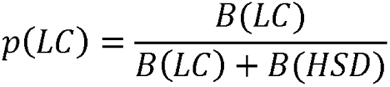

and thus 1-*p(LC)* can be viewed as the probability that the HSD-model (null-model) is a better model.

### Construction and validation of the mRNA expression CELTYC classifier (LIHC)

We built an mRNA-expression based predictor to classify mRNA expression profiles into one of the two CELTYC prognostic subgroups, defined as the 2 main clusters obtained using lymphocyte specific DMCTs, i.e. we assigned a sample to good-outcome (Gclust) if it was part of LC2, or poor outcome (Pclust) if it was part of LC1/LC3. Using the assignment of the 373 LIHC TCGA tumor samples into poor- and good-outcome groups encoded as 1 and 0 respectively, we trained a logistic lasso model with *glmnet* as follows. First, the expression profiles for each gene were scaled to unit variance. To select the optimal penalty parameter (lambda) value, we implemented a nested 10-fold cross-validation ^115^. This procedure involved splitting the samples into 10 folds. Using 9 folds, a series of logistic lasso models with 500 lambda values ranging from 0 to 0.1 were trained and finally tested on the leave-out bag. This was repeated 9 times so that each fold was used for testing exactly once. For each fold, we recorded the predicted values and for each choice of the 500 different lambda values. For each lambda value, we then concatenated the scores (predicted probabilities) of all folds, and computed an AUC value with R package *pROC.* The lambda value with the greatest AUC was selected as the optimal parameter to retrain a final logistic lasso model using all TCGA LIHC samples.

### Validation of the CELTYC lasso mRNA expression predictor

We validated the lasso predictor of clinical outcome in two independent LIHC mRNA expression datasets ^49,50^. Normalized expression data for 100 HCC samples (GSE16757) and 246 HCC samples (GSE14520) were downloaded from GEO. The GSE16757 expression data was processed on Illumina BeadStudio software and normalized using quantile normalization and log2 transformation, while the GSE14520 expression data containing samples from 2 cohorts was processed with the *matchprobes* package and the RMA method in the R *affy* package, followed by log2-transformation. Gene expression profiles for genes in our lasso predictor were scaled to unit variance. Using the estimated regression coefficients beta from the optimal lasso logistic model, we then calculated the linear predictor scores (LP), i.e. X*beta+intercept, and finally transformed the scores to probabilities of belonging to poor outcome class, using exp(LP)/(1+exp(LP)). Cox regressions were performed correlating the predicted probabilities to overall survival. For the Kaplan Meier analysis, we defined samples with the upper and lower 25% quantile probability scores as “poor outcome” and “good outcome”, respectively. We also performed Cox regressions after merging the predicted probabilities of both datasets and merging the predicted groups of both datasets.

### Somatic mutational and copy-number variation data of TCGA LIHC tumors

Masked somatic mutation data for LIHC was downloaded with *TCGAbiolinks* Bioconductor R-package. For each gene, samples with any of the following mutation-types (missense mutation, silent mutation, frameshift deletion, in-frame deletion, frameshift insertion, intron mutation, 3’ UTR mutation, splice-site mutation, nonsense mutation, and splice-region mutation) were encoded as 1, otherwise encoded as 0. Fisher exact tests were then performed to identify genes with significantly different mutation frequency between the 2 CELTYC subgroups of different prognosis. We identified the survival associated genes from the ones with different mutation frequency between CELTYC subgroups by doing cox regressions of mutational profiles against survival, and incorporated the survival associated genes with CELTYC subgroups in a multivariate cox regression to see whether the CELTYC classification is independent of mutational profiles.

Gene level copy number data of around 56k genes for TCGA LIHC was downloaded with *TCGAbiolinks* package. For each gene, samples with loss or deletions (i.e. copy number < 2) were encoded as 1, otherwise encoded as 0 (i.e. copy number >=2) and Fisher exact tests were performed to see whether the deletion/loss incidence for each gene is significantly different between the two CELTYC subgroups. An analogous analysis was done for gain/amplification.

### Construction and validation of the DNAm CELTYC Epi and IC classifiers (KIRC)

Clustering over the epithelial specific DMCTs resulted in an assignment of KIRC samples into one of 3 clusters, labeled by an ordinal integer (1,2,3) with 3 and 1 labeling the poor and good outcome clusters, respectively. A similar clustering-scheme was obtained for the immune-cell specific DMCTs. We then used an elastic net (alpha=0.5) classifier (glmnet R-package) with a 5-fold cross-validation strategy to build a DNAm-predictor of the cluster labels, treating it as an ordinal variable and optimizing the root-mean-square error (RMSE). The training and testing of the predictor was done on the residual matrix obtained after regressing out the effect of cell-type fractions and defined over the respective cell-type specific DMCTs. Of note, since these DMCTs were inferred by comparing cancer to normal tissue, it is legitimate to then train the predictors of CELTYC clusters over these DMCTs, since this training is only done over cancer samples. When implementing the 5-fold CV, we ensured that folds had proportional numbers of cancer samples belonging to each cluster. Optimal penalty parameter was tuned on the combined left-out sets. For validation, we used an independent Illumina 450k DNAm dataset of 115 ccRCC samples ^87^. Before applying the Epi and IC-predictors, we estimated epithelial, fibroblast and immune-cell fractions in the independent dataset, generating residuals after regressing out the effect of these cell-type fractions. The scores from the Epi and IC-predictors were then assigned to 3 clusters by ranking the scores and using the same quantiles inferred from the training set.

### Cell-proliferation index and mitotic-age

The cell proliferation index was computed from the RNA-Seq data of the LIHC samples and measures the instantaneous rate of cell proliferation of the tumor. It was computed using a set of cell proliferation genes by z-scoring their values and then averaging over them, as described by us previously ^116^. Mitotic-age of a tumor sample is an estimate of the total cumulative number of stem-cell divisions of a tissue and is computed from DNAm data. Here we applied our epiTOC2 mitotic age clock ^83^ to the KIRC samples. epiTOC2 yields estimates for both the total number of stem-cell divisions (TNSC) (age-dependent) and the average lifetime intrinsic rate of stem-cell division (irS) (which is naturally age-adjusted).

### Overrepresentation and Gene Set Enrichment Analyses

*LIHC:* To test for enrichment overrepresentation of cell-type specific hypermethylated and hypomethylated DMCTs in LIHC, we used the web-based tool eFORGE ^54^. We also performed GSEA ^55,56^ for genes ranked by upregulation in the poor outcome CELTYC cluster using the *clusterProfiler* R-package ^117^ focusing on the MsigDB Hallmark gene set. With bulk RNA-Seq data and the barcodes which could be matched to methylation data, DEGs were identified between Pclust and Gclust by applying Limma on bulk RNA-Seq data, adjusting for age, sex and cell-type fractions. The DAVID webtool ^118,119^ was run on genes whose expression correlated with poor outcome according to Cox-regression analyses adjusted for cell-type fractions.

*KIRC:* We performed enrichment overrepresentation analysis focusing on genes containing epithelial, fibroblast and immune-cell specific cancer-DMCTs, stratified according to up or downregulation in cancer vs normal tissue (adjusted for cell-type fractions). Only genes with FDR<0.05 were selected. If number was larger than 250, the top-250 were used. Overrepresentation analysis was performed using a one-tailed Fisher’s exact test, as implemented in ebGSEA ^80^. We also performed ebGSEA on the top-250 genes whose expression correlates positively and negatively with the CELTYC clusters, adjusted for cell-type fractions. Both overrepresentation analyses were done using the Cancer Hallmark gene set collection from MSigDB ^56^.

*BRCA:* We performed enrichment overrepresentation analysis using ebGSEA and MSigDB as described for KIRC, using the cancer hallmark, cell-type and immune-cell signature collections.

### Cytokine activity scores

We used a large compendium of cytokine stimulation signatures from the Immune Dictionary ^70^. Briefly, these are perturbation mRNA signatures obtained from scRNA-Seq data, where specific immune cell-types were stimulated with a variety of cytokines. A total of 938 cytokine-response signatures are available encompassing 17 immune cell types with a mean of 55 cytokines tested for each cell-type. Cytokine signatures were first filtered by the requirement that they contain at least 10 upregulated genes with representation in the TCGA LIHC and KIRC RNA-Seq datasets. Of note, the genes in the RNA-Seq datasets were first z-score normalized, hence we required that genes be expressed in at least 10% of all samples, to avoid singularities/outliers with zero or ultra-low variance. The requirement that cytokine signatures should have at least 10 upregulated genes, resulted in reduced sets of 342 and 333 cytokine signatures for the LIHC and KIRC RNA-Seq datasets, respectively. Subsequently, a cytokine activity score for each sample was computed by averaging the z-score normalized expression of all upregulated cytokine signature genes. This score was then correlated to CELTYC clusters or clinical outcome.

### Identification of cell-type specific cancer associated differentially expressed genes

We applied CIBERSORTx^76^ via the CIBERSORTx webtool (https://cibersortx.stanford.edu/) on the LIHC expression data to identify cell-type specific differentially expressed genes. The gene expression reference matrix defined over 658 marker genes for 5 cell types (cholangiocytes, endothelial, hepatocytes, Kupffer cells, lymphocytes) obtained from R package EpiSCORE was input as the “signature matrix” for CIBERSORTx. CIBERSORTx group-mode imputes average values of cell type-specific gene expression profiles for each gene in each individual cell type from a group of bulk tissue expression profiles, and outputs the corresponding standard error values for each gene. Therefore, to identify cell-type specific cancer associated DEGs, we ran group-mode CIBERSORTx with expression data for LIHC cancer samples and normal samples separately. Z-statistics and P values were then calculated using the output average expression values and standard errors for the two groups to identify significant DEGs.

### Combinatorial clustering/indexing and prognostic synergy in KIRC

Given the clusters inferred from cell-type specific DMCTs, combinatorial clustering or indexing refers to the construction of new clusters made up of the various combinations of cell-type specific clusters. For instance, for two cell-types, each predicting 3 clusters, combinatorial clustering leads to 9 clusters. To formally test that a Cox-regression model against the 9 clusters (categorical variable, 8 degrees of freedom) is a better prognostic model than the ones defined by each of the cell-type specific clusters (2 degrees of freedom), we used a likelihood ratio test (LRT) i.e. a one-tailed Chi-square test with 6 degrees of freedom. For the case where we treat specific clusters in the combinatorial model as ordinal (thus a model with only 1 degree of freedom), we can’t use a LRT test, because each of the cell-type specific clusters, if treated as ordinal, also define models with only 1 degree of freedom. Hence, in this case we derive relative probabilities for the respective models, as follows: First, we compute the Akaike Information Criterion (AIC) values ^114^ for each model *m:*

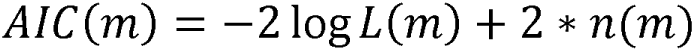

where *L(m)* is the model likelihood and *n(m)* is the number of parameters of model *m.* Since in the ordinal case, all models have the same number of parameters to be inferred, we can ignore this term. Then, for two models *k* and *j* we define

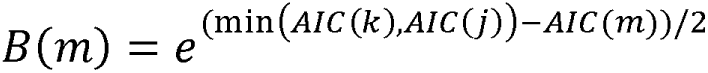

where *m* can be either *k* or *j.* Finally, the relative probability of the two models being true is given by

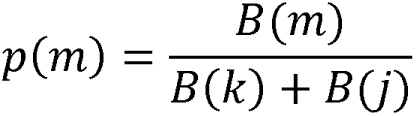

For instance, for the combinatorial ordinal cluster model (cmb), its relative probability of being true relative to the model defined by ordinal immune-cell (IC) clusters, would be:

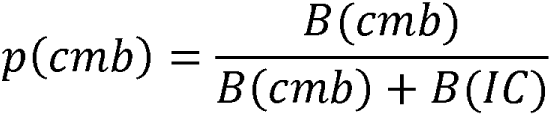

and so *1*-*p(cmb)* can be viewed as the probability that the IC-model (null-model) is a better model.

### Prognostic macrophage signature evaluation in KIRC

For the prognostic macrophage signatures ^84^ we computed the scores by first z-scoring the mRNA expression KIRC dataset, and then averaging the corresponding genes. For the chemokine/cytokine signature, we averaged z-scores of *CXCL8, CXCL2, CCL4, CCL3, CCL4L2, CXCL3, CCL3L3, CCL20, NFKB1* and *IL1B*. For the lysosomal signature, we averaged z-scores of *CTSL, ASAH1, LGMN, LIPA, CTSD* and *LAMP1*. These scores were then correlated to overall survival with Cox-regression and the CELTYC ordinal clusters with a linear regression.

### CELTYC application to RRBS breast cancer DNAm dataset

We downloaded the normalized RRBS MetaBric dataset ^88^, with DNAm summarized at the gene promoter level. After mapping to Entrez gene IDs, we were left with a DNAm data matrix defined for 12,338 genes and 1710 samples. We further restricted the analysis to 231 normal-adjacent and 797 luminal ER+ cancer samples. Cell-type fractions for 7 cell-types (Fibroblasts, Fat, Endothelial Cells, Lymphocytes, Macrophage, luminal and basal epithelial cells) were estimated using EpiSCORE’s DNAm reference matrix for breast. CellDMC was run comparing normal-adjacent to the 797 luminal ER+ samples, and CELTYC subsequently applied to lymphocyte, endothelial and luminal differentially methylated gene promoters. Mitotic age was estimated using epiTOC2 ^83^. Survival analysis was performed with survival R-package.

## Code Availability

R-functions for running the CELTYC algorithm are available from https://github.com/QL2024/CELTYC.

## Data Availability

The Illumina DNA methylation datasets used here are freely available from the public repository GEO (www.ncbi.nlm.nih.gov/geo) under the following accession numbers: GSE213478 (eGTEX lung), GSE40279 (whole blood), GSE210843 (bronchial epithelial cells) and GSE113501 (ccRCC + kidney controls). The two LIHC RNA-Seq datasets are freely available on GEO under accession numbers GSE14520 and GSE16757. TCGA Illumina 450k datasets for LIHC and KIRC are available from the GDC data portal https://portal.gdc.cancer.gov/. The BLUEPRINT DNA dataset of sorted immune cell-types is available from EGA https://ega-archive.org/ under accession number EGAS00001001456. The breast cancer MetaBric RRBS dataset is available from https://tanaylab.weizmann.ac.il/metabric_rrbs. The liver, kidney and breast DNAm-reference matrices are freely available from the EpiSCORE and EpiDISH R-packages https://github.com/aet21/EpiSCORE and https://bioconductor.org/packages/release/bioc/html/EpiDISH.html

## Declarations

### Ethics approval and consent to participate

Not applicable here since we have only analyzed existing publicly available data.

### Consent for publication

Not applicable.

### Availability of data and materials

The Illumina DNA methylation datasets used here are freely available from the public repository GEO (www.ncbi.nlm.nih.gov/geo) under the following accession numbers: GSE213478 (eGTEX lung), GSE40279 (whole blood), GSE210843 (bronchial epithelial cells) and GSE113501 (ccRCC + kidney controls). The two LIHC RNA-Seq datasets are freely available on GEO under accession numbers GSE14520 and GSE16757. TCGA Illumina 450k datasets for LIHC and KIRC are available from the GDC data portal https://portal.gdc.cancer.gov/. The BLUEPRINT DNA dataset of sorted immune cell-types is available from EGA https://ega-archive.org/ under accession number EGAS00001001456. The breast cancer MetaBric RRBS dataset is available from https://tanaylab.weizmann.ac.il/metabric_rrbs. The liver, kidney and breast DNAm-reference matrices are freely available from the EpiSCORE and EpiDISH R-packages https://github.com/aet21/EpiSCORE and https://bioconductor.org/packages/release/bioc/html/EpiDISH.html. R-functions for running the CELTYC algorithm are available from https://github.com/QL2024/CELTYC.

### Competing interests

The authors declare no competing interests.

### Funding

This work was supported by NSFC (National Science Foundation of China) grants, grant numbers 32170652, 31970632 and 32370699.

### Authors contributions

LQ and AET performed the statistical analyses. AET conceived and designed the study. AET wrote the manuscript with contributions from LQ.

## Acknowledgements

We would like to thank the TCGA and everyone who supports open-access data.

## Notes

### Competing Interest Statement

The authors have declared no competing interest.

